# Genome-wide SNP genotyping of DNA pools identifies untapped landraces and genomic regions that could enrich the maize breeding pool

**DOI:** 10.1101/2020.09.30.321018

**Authors:** Mariangela Arca, Brigitte Gouesnard, Tristan Mary-Huard, Marie-Christine Le Paslier, Cyril Bauland, Valérie Combes, Delphine Madur, Alain Charcosset, Stéphane D. Nicolas

**Affiliations:** Université Paris-Saclay, INRAE, CNRS, AgroParisTech, GQE - Le Moulon, 91190, Gif-sur-Yvette, France; Amélioration Génétique et Adaptation des Plantes méditerranéennes et tropicales, Univ Montpellier, CIRAD, INRAE, Institut Agro, F-34090 Montpellier, France; Université Paris-Saclay, INRAE, Etude du Polymorphisme des Génomes Végétaux, 91000, Evry, France

**Keywords:** *Zea mays*, gene banks, Landraces, Pre-breeding, DNA pooling, Genetic diversity, Selective footprints, Allelotyping

## Abstract

Maize landraces preserved in genebanks have a large genetic diversity that is still poorly characterized and underexploited in modern breeding programs. Here, we genotyped DNA pools from 156 American and European landraces with a 50K SNP Illumina array to study the effect of both human selection and environmental adaptation on the genome-wide diversity of maize landraces. Genomic diversity of landraces varied strongly in different parts of the genome and with geographic origin. We detected selective footprints between landraces of different geographic origin in genes involved in the starch pathway (*Su1, Waxy1*), flowering time (*Zcn8, Vgt3, ZmCCT9*) and tolerance to abiotic and biotic stress (*ZmASR, NAC* and *dkg* genes). Landrace diversity was compared to that of (i) 327 inbred lines representing American and European diversity (“CK lines) and (ii) 103 new lines derived directly from landraces (“DH-SSD lines”). We observed limited diversity loss or selective sweep between landraces and CK lines, except in peri-centromeric regions. However, analysis of modified Roger’s distance between landraces and the CK lines showed that most landraces were not closely related to CK lines. Assignment of CK lines to landraces using supervised analysis showed that only a few landraces, such as Reid’s Yellow Dent, Lancaster Surecrop and Lacaune, strongly contributed to modern European and American breeding pools. Haplotype diversity of CK lines was more enriched by DH-SSD lines that derived from the landraces with no related lines and the lowest contribution to CK lines. Our approach opens an avenue for the identification of promising landraces for pre-breeding.

**SIGNIFICANCE STATEMENTS:** Maize landraces are a valuable source of genetic diversity for addressing the challenges of climate change and the requirements of low input agriculture as they have been long selected to be well adapted to local agro-climatic conditions and human uses. However, they are underutilized in modern breeding programs because they are poorly characterized, genetically heterogeneous and exhibit poor agronomic performance compared to elite hybrid material. In this study, we developed a high-throughput approach to identify landraces that could potentially enlarge the genetic diversity of modern breeding pools. We genotyped DNA pools from landraces using 50K array technology, which is widely used by breeders to characterize the genetic diversity of inbred lines. To identify landraces that could enrich the modern maize germplasm, we estimated their contribution to inbred lines using supervised analysis and a new measurement of genetic distance.

## INTRODUCTION

Plant genetic resources are the basic raw material for future genetic progress (1–4). Maize landraces are an interesting source of genetic diversity for addressing the challenges of climate change and the requirements of low input agriculture, as they have been long selected to be well adapted to local agro-climatic conditions and human uses (4–7). During the early twentieth century, landraces were used as parent material for the development of improved hybrid varieties to meet the needs of modern agriculture. During this transition from landraces to hybrids, many favorable alleles were probably lost as a result of their association with unfavorable alleles and/or genetic drift (8–11). Nowadays, modern breeding programs tend to focus on breeding populations that can be traced back to a few ancestral inbred lines derived from landraces at the start of the hybrid era (12–15). Landraces that did not contribute to this founding material may be expected to be useful for enriching modern maize diversity, particularly for traits that enhance adaptation to adverse environmental conditions (7). However, maize landraces are used to a very limited extent, if at all, in modern plant breeding programs because they are poorly characterized, genetically heterogeneous and exhibit poor agronomic performance compared to elite hybrid material (3, 6, 16–18). Therefore, understanding the genetic diversity of maize landraces and their relation to the maize elite pool is essential for better management of genetic resources and for genetic improvement through genome-wide association studies, genomic selection and the dissection of quantitative traits (2, 6, 7).

Maize was domesticated in the highlands of Central Mexico approximately 9,000 years ago (19, 20). It then diffused to South and North America (21, 22) and spread rapidly out from America (23). It is now cultivated in highly diverse climate zones ranging from 40°S to 50°N. In Europe, the presently accepted hypothesis is that maize was first introduced through Spain by Columbus, although other sources of maize that were pre-adapted to temperate climates have been important for adapting to northern European conditions (22, 24–29). After being introduced in different parts of the world, maize landraces were then selected by farmers to improve their adaptation to specific environments, leading to changes in flowering behavior, yield, nutritive value and resistance to biotic and abiotic stress, resulting in subsequent differentiation of the material (7, 27, 30).

In recent years, the genetic diversity of maize landraces, which are conserved *ex situ*, has been studied extensively using various types of molecular markers such as restriction fragment length polymorphisms (RFLPs) (8, 25–28, 31–34) and simple sequence repeats (SSRs) (8, 23, 35, 36). Single nucleotide polymorphisms (SNPs) are now the marker of choice for various crop species such as maize (37), rice (38) and barley (39). They are the most abundant class of sequence variation in the genome, are co-dominantly inherited, genetically stable, easily automated and, thus, suitable for high-throughput automated analysis (40). Unlike SSRs, allele coding can be easily standardized across laboratories and the cost of genotyping is very low, which is a major advantage for characterizing genetic resources. A maize array with approx. 50,000 SNP markers has been available since 2010 (37). It has been successfully used to analyze the diversity of inbred lines and landraces by genotyping a low number of plants per accession (13, 16, 41–44).

However, due to high within-accession diversity, the characterization of maize landraces should be carried out on a representative set of individuals (45). Despite recent technical advances, genotyping large numbers of individuals remains very expensive in the context of genetic characterization. As a result, DNA pooling (or allelotyping) has been actively developed as a valuable alternative strategy for collecting information on allele frequency from a group of individuals while significantly reducing the effort required for population studies using DNA markers (46, 47). In maize, DNA pooling has been successfully used to decipher the global genetic diversity of landraces using RFLP (32) and SSR markers (23, 27, 28, 48, 49). The recent development of SNP arrays in maize (37, 50), combined with DNA pooling, could be useful for characterizing the genetic diversity of maize landraces at a fine genomic scale. In a previous study, we developed a new method for predicting the allelic frequency of each SNP from a maize Illumina 50K array within DNA pools based on the fluorescence intensity of the two alleles at each SNP (51). This new method accurately predicts allelic frequency, safeguards against the false detection of alleles and leads to little ascertainment bias for deciphering global genetic diversity (51).

In the present study, we applied this new method on a pilot scale to: i) investigate the genome-wide diversity and genetic structure of 156 maize landraces that are representative of European and American diversity; ii) compare the diversity of these landraces to that of a panel of 327 inbred lines that represent the diversity presently used in North-American and European breeding, the “CK lines” (27) and 103 new inbred lines derived from landraces, the “DH-SSD lines”; and iii) identify the landraces that could potentially broaden the genetic diversity of the CK lines.

## RESULTS

### Genetic diversity within maize landraces

Only 25 SNPs out of 23,412 were monomorphic in the landrace panel. The average total diversity (HT) was 0.338 ± 0.001. The distribution of minor allelic frequency (MAF) showed a deficit in rare alleles (MAF<0.05) compared to other frequency classes (Fig. S1).

In order to compare the genetic diversity of populations from different regions, we classified the 156 landraces into five geographic groups: Europe (EUR), North America (NAM), Central America and Mexico (CAM), the Caribbean (CAR) and South America (SAM) (Table 1, Fig. S2, Table S1). All five geographic groups displayed both alleles for nearly all loci, with the exception of CAR which was monomorphic at 1,227 loci out of 23,387 (Fig. S3). The lowest and highest within-group HT was found in CAR (0.301) and CAM (0.328), respectively. Note that there was an excess of rare alleles in EUR, CAR and NAM but not in SAM and CAM (Fig. S1).

**Table 1:**
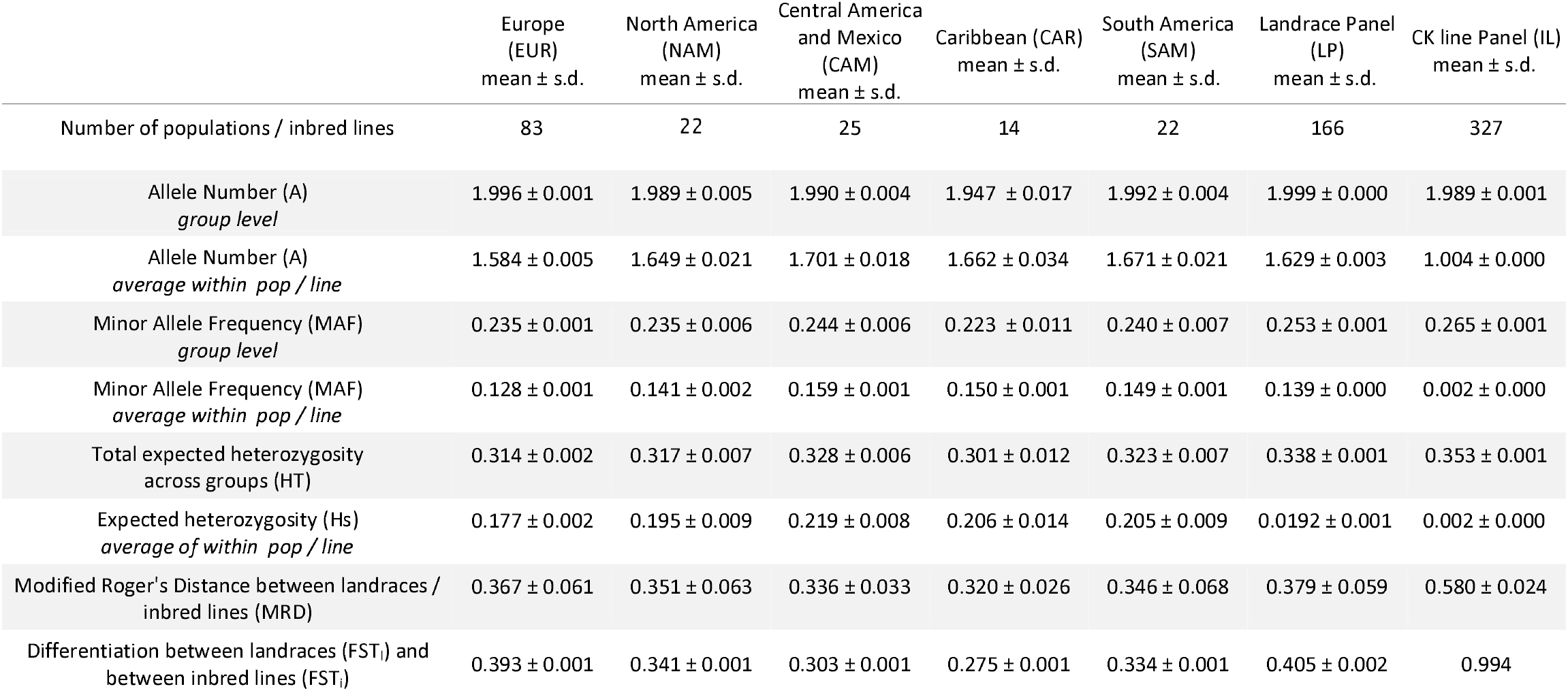
Genetic diversity within the five geographic groups of landraces, the entire landrace panel and the CK line panel.

The average number of alleles per locus and per landrace within the entire landrace panel was 1.629 ± 0.003 and ranged from 1.098 (Ger8) to 1.882 (Sp11). Gene diversity within landraces (Hs) was on average 0.192 ± 0.001, (Table 1) and varied between 0.03 (Ger8 and Ger9) and 0.28 (Sp11) (Table S1). The CAM group displayed on average the highest diversity (0.219 ± 0.008), while the EUR group displayed the lowest (0.177 ± 0.002).

Genetic differentiation between landraces (FST) was 0.428 on average. FST within a geographic group varied between 0.314 (CAR) and 0.434 (EUR) (Table 1). Overall genetic differentiation between geographic groups was low (FST=0.05). FST between pairs of geographic groups varied between 0.016 (EUR and NAM) and 0.083 (NAM and CAR) (Table S2).

### Relationship between maize landraces and population structure

The average modified Roger’s distance (MRD) between landraces was 0.379. The lowest MRD between landraces was 0.158 (Chi12 and Chi9). It is slightly higher than the distance between two pools of independent individuals from a same population (0.092-0.120, (51)). The highest MRD was 0.552 (Ant1 and Ger8). The average MRD between populations from a same geographic group ranged from 0.320 (CAR) to 0.367 (EUR) (Table 1). The average MRD between populations belonging to two different geographic groups varied between 0.354 (CAM *vs* CAR) and 0.420 (NAM *vs* CAR) (Table S2).

We investigated the relationship between maize landraces using Principal Coordinate Analysis (PcoA) and Ward hierarchical clustering based on MRD (Fig. 1). For both PcoA and Ward hierarchical clustering analysis, landraces mostly clustered according to their geographic proximity (Fig. 1, Fig. S4, Fig. S5). The first axis (PC1, 18.4% of the total variation) discriminated (i) temperate landraces belonging to the Northern Flint cluster (from northern Europe and North America) from (ii) tropical and subtropical landraces (from the Caribbean and South and Central America) (Fig. 1A). The second axis (PC2, 5% of the total variation) discriminated (i) North American (Corn Belt Dent cluster), Central American and Mexican populations (Mexican cluster) from (ii) Italian (Italian Flint cluster), and Spanish and French populations (Pyrenean-Galician cluster). Ward hierarchical clustering showed that at the highest level (k=2, Fig. 1B), 62 of the 83 European landraces clustered together (European cluster) while 70 of the 83 American landraces clustered together (American cluster). At a deeper level (k=7), we distinguished 4 clusters of American or European landraces, each originating from a geographic area with homogeneous agro-climatic conditions (cluster a, b, e and f in Fig. 1B, Fig. S4). Cluster “a” grouped 15 landraces that originated mainly in Mexico and southwestern USA. Cluster “b” comprised 10 South American landraces that originated along the Andean Mountains. Cluster “e” grouped 31 European landraces that originated either along the Pyrenean Mountains or in Central Eastern Europe. Cluster “f” grouped mainly Italian Flint landraces. Three clusters grouped together American and European landraces (cluster c, d and g on Fig. S4). Cluster “c” comprised 14 dent landraces that originated mainly from Eastern European landraces and the US Corn Belt. Cluster “d” grouped 65 landraces mostly from southern Spain (latitude <40°N), southwestern France and from the Caribbean Islands and countries bordering the Caribbean Sea (d1, d2 and d3 on Fig. S4). Cluster “g” comprised 12 North American flint landraces from higher latitudes (>40°N) and 18 northeastern European landraces mainly from Germany (g on Fig. S4). Using a pairwise Mantel test for each geographic area, we observed a low but significant correlation between the genetic distance and geographic distance matrices for EUR (r^2^ = 0.05, P < 0.001, Fig. S6A), NAM (r^2^ = 0.12, P < 0.001, Fig. S6B) and CAM (r2 = 0.0858, P = 0.02, Fig. S6C).

**Fig. 1:**
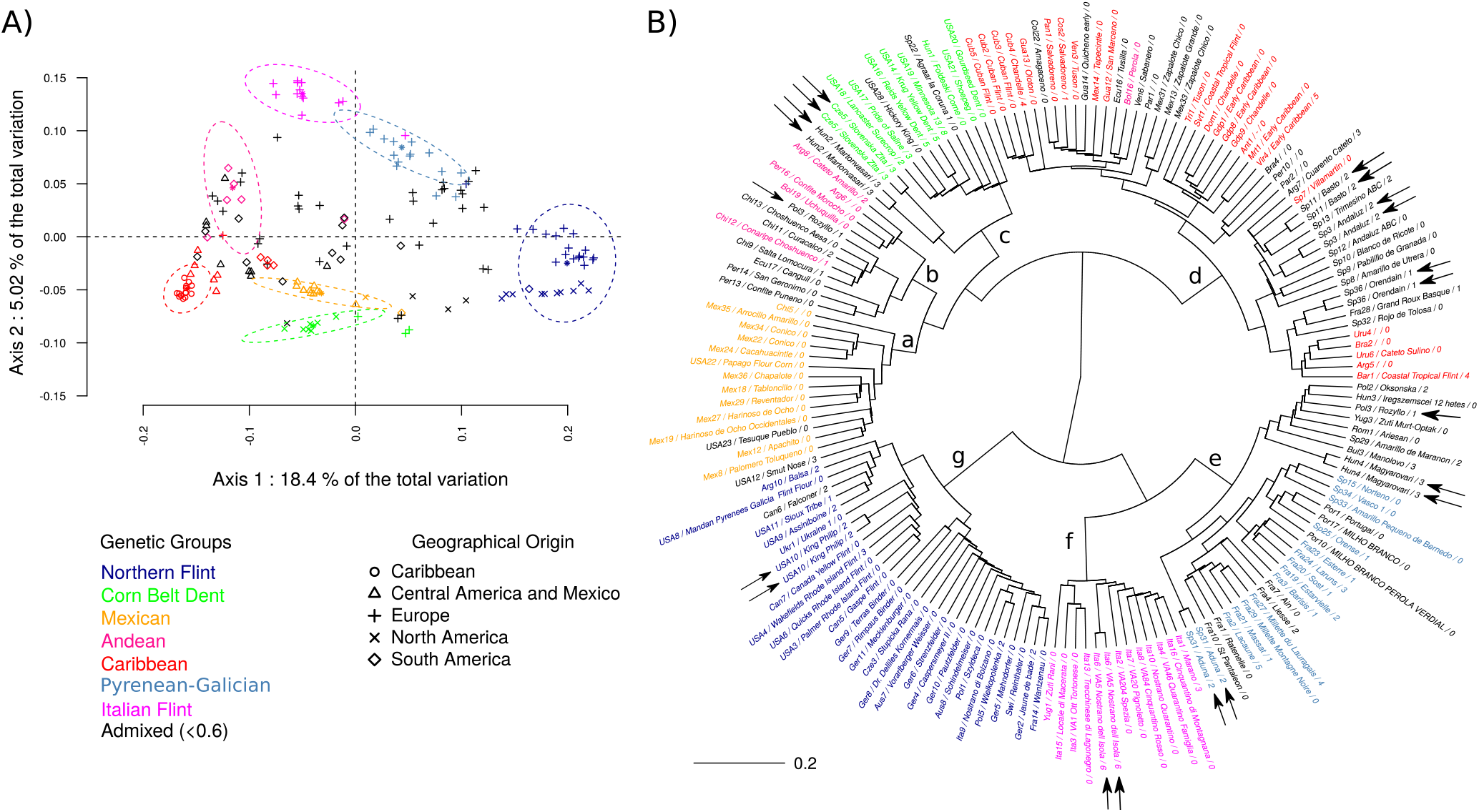
Genetic relationship between 156 maize landraces based on their modified Roger’s distance (MRD). A) Projection of the 166 DNA samples on the first two axes of the Principal Coordinate Analysis. Symbols indicate the geographic origin of landraces. B) Dendrogram obtained by Hierarchical clustering, using Ward’s algorithm. Labels indicate for each landrace their abbreviation code, common names and number of first cycle inbred lines they contributed to, respectively. Black arrows indicate the 10 landraces with duplicated DNA samples. Colors indicate the assignment of landraces to the seven genetic groups defined by ADMIXTURE. Landraces with an assignment probability below 0.6 were considered admixed and colored in black.

We analyzed the genetic structure of 156 landraces using the ADMIXTURE program. Likelihood analysis indicated that the optimal number of genetic groupe was K=2, K=3 and K=7 (Fig. S7). We considered K=7 as the reference, as this value was consistent with the one obtained with 24 SSRs by Camus-Kulandaivelu et *al*. (27). Landraces from different geographic regions were assigned to different genetic groups, with a clear trend along latitude and longitude. Fig. 2). Assignment to these groups was also highly consistent with PcoA and hierarchical clustering (Fig. 1, Fig. 2, Fig. S4, Fig. S5). The genetic structure obtained with SNP markers was highly consistent with that obtained with the 17 SSR markers; indeed, 72% (K=7) to 100% (K=3) of landraces were assigned to the same group by both types of markers (Table S3). The main differences between the SSR and SNP results at K=7 were that the Northern Flint landrace group obtained with SNPs is split in two with SSRs and the separate Pyrenean-Galician and Italian groups found with SNPs form a single group with SSRs.

**Fig. 2:**
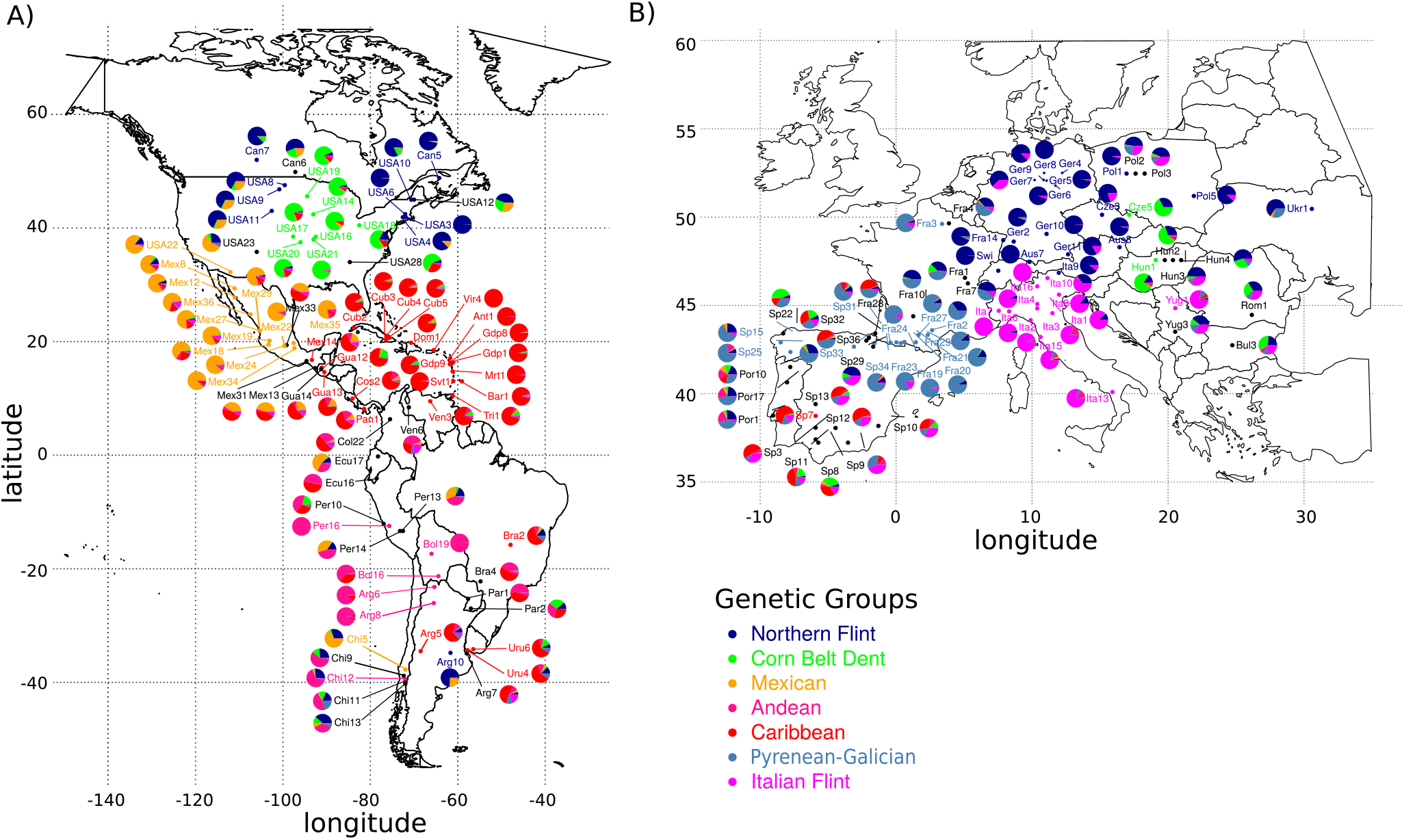
Spatial genetic structure of American (A) and European (B) maize landraces. Population structure is based on ADMIXTURE analysis with K = 7. Each population is represented by a pie diagram whose composition indicates admixture coefficients. Population labels are colored according to their main assignment (>0.6), and are black if the landrace is admixed.

### Scanning the maize landrace genomes for regions under selection

Using a sliding window approach, we identified 14 regions with windows containing at least two SNPs with extremely low genetic diversity 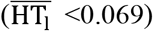 across the entire landrace panel (Fig. 3A, Table S4). These regions were mainly located in the centromeric region of chromosomes 5 and 7. Genomic regions showing low diversity within geographic groups were most abundant in CAR (67), followed by EUR (56), CAM (39), SAM (36) and NAM (26) (Fig. 3E to 3I, Table S4). These regions were mostly located close to the centromeres but varied between geographic groups. In the centromeric region of chromosome 1, we observed (i) no loss of diversity for CAR and NAM and (ii) a depletion in genetic diversity for CAM, EUR and SAM. Conversely, we observed a strong depletion on chromosomes 3 and 4 in CAR landraces that was not observed in other geographic groups.

**Fig. 3:**
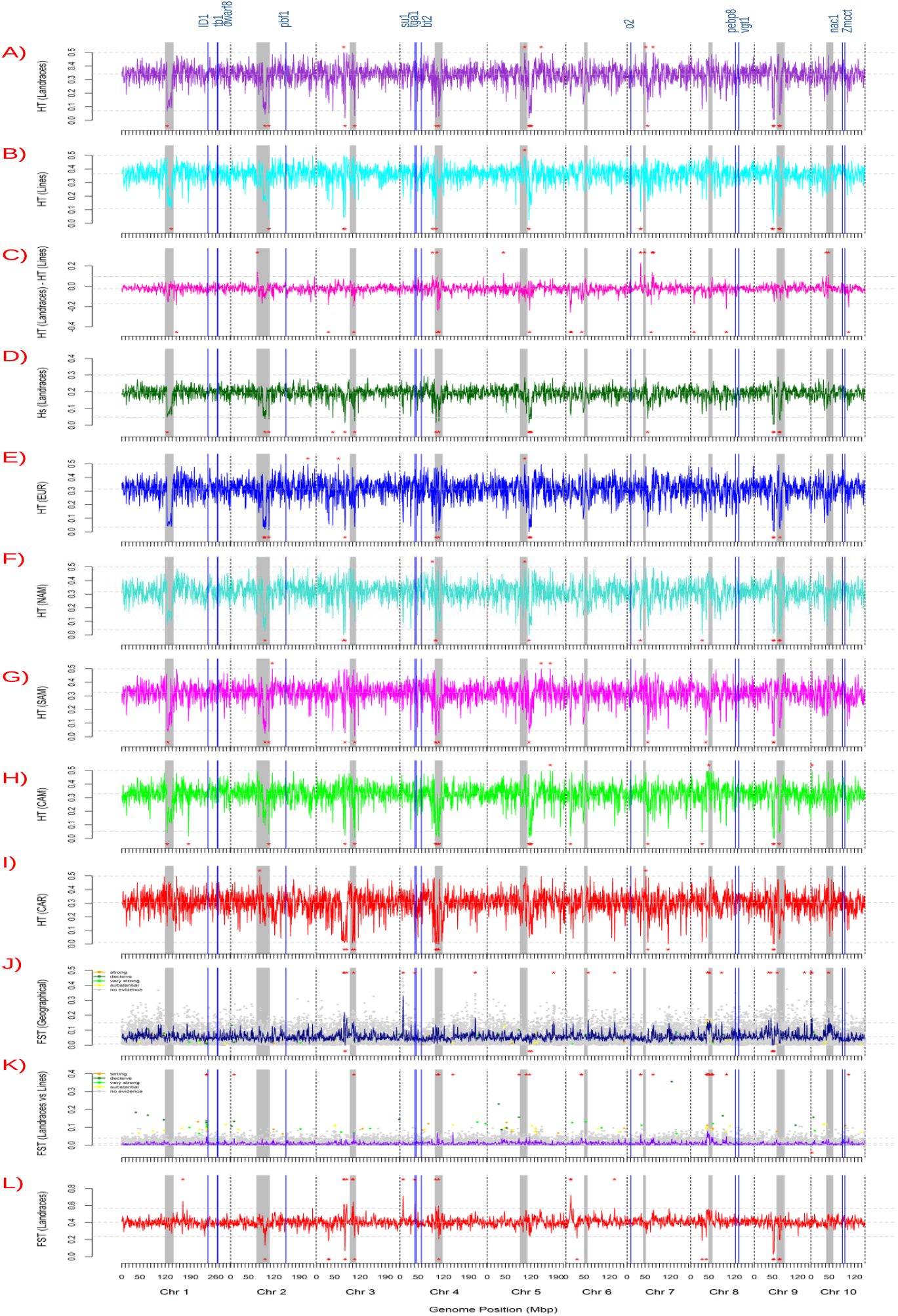
Variation in genetic diversity and differentiation along the maize genome. A) Total expected heterozygosity across landraces: HT (Landraces); B) total expected heterozygosity (HT) across inbred lines: HT (Lines); C) difference between the total expected heterozygosity across landraces and across inbred lines: HT (Landraces) – HT (Lines); D) mean expected heterozygosity within landraces: Hs (Landraces); total expected heterozygosity across landraces from E) Europe: HT (EUR)), F) North America: HT (NAM), G) South America: HT (SAM); H) Central America and Mexico: HT (CAM), I) the Caribbean: HT (CAR), J) FST between geographic groups of landraces: FST (Geographic); K) FST between landraces and inbred lines: FST (Landraces vs. Inbred lines); L) FST between landraces: FST (Landraces). Loci with decisive, very strong, strong, substantial, no evidence of selection using bayescan are colored in orange, dark green, light green, yellow and blue (J, K, L). Vertical gray bars correspond to centromere limits. Chromosome boundaries are indicated by vertical dashed lines. Horizontal dashed lines correspond to the mean, 5^th^ and 95^th^ percentile of each parameter. Outlier regions are indicated by red asterisks (>95% at the top, <5% at the bottom). Vertical blue lines indicate the location of the genes *ID1, tb1, pbf1, su1, tga1, bt2, o2, pebp8, vgt1, nac1* and *Zmcc*

Outlier analysis of FST values among individual landraces identified 20 and 17 genomic regions displaying high differentiation 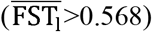 and low differentiation 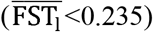 between landraces, respectively (Fig. 3L, Table S4). Genetic differentiation was highest upstream of chromosome 6 (Sp10 in Table S5), in two regions upstream of chromosome 4 (Sp6 and Sp7in Table 2) and in one region on chromosome 3 (Sp3 in Table 2).

**Table 2:**
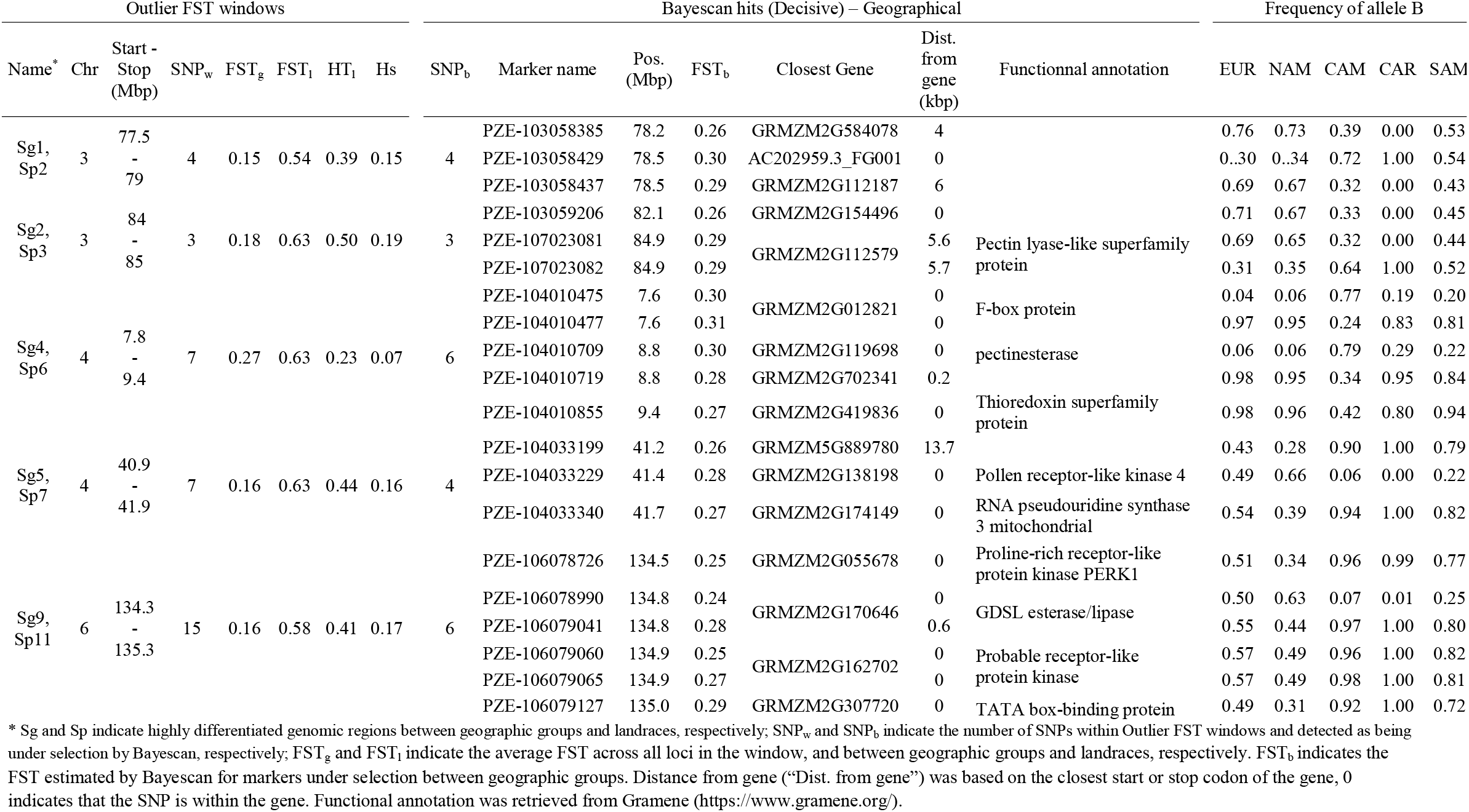
Genomic regions identified as being highly differentiated between landraces and geographic groups. Only SNPs that were detected by BAYESCAN with decisive evidence of selection and Outlier FST windows carrying at least two SNPs are listed.

Outlier FST analysis between geographic groups identified 26 regions with high differentiation 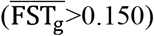 and 8 regions with low differentiation 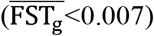 (Fig. 3J, Table S4); BAYESCAN identified 379 loci under divergent selection (Fig. 3J, Table S6 and S7, Fig. S8). The five genomic regions that were previously identified as being highly differentiated between landraces by outlier FST analysis were also detected by both FST outlier and BAYESCAN analyses between geographic groups. (Table 2). Only one highly differentiated genomic region was identified between landraces but not between all five geographic groups (Sp10 in Table S5) whereas 6 genomic regions were identified exclusively between the five geographic groups (Sg6, 7, 13, 15, 17, 18 and 20 in Table S5). These regions displayed contrasted allelic patterns across geographic groups. Sp10 (11.7 Mbp – 15.3 Mbp on chromosome 6, 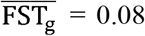 and 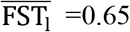) had 9 SNPs that were close to fixation in CAM (HT<0.1), but were segregating in NAM (~0.4) and also to a lesser extent in EUR, CAR and SAM (HT~0.2). Sg2-Sp3 (84-85 Mbp on chromosome 3, 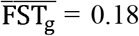 and 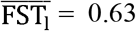) had 3 SNPs showing a continuous allelic frequency gradient between tropical and temperate landraces with one allele largely predominant in NAM and EU (~70%), minor in CAM (~30%) and absent in CAR (~0%). Sg4-Sp6 (40.3-41.8Mbp on chromosome 4, 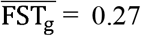 and 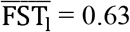) had 4 SNPs that were nearly fixed in temperate landraces (NAM, EUR) and displaying intermediate frequencies in CAM. By contrast, the Sg5-Sp7 region 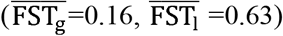 displayed higher diversity in temperate (HT_NAM_ and HT_EUR_~0.4) than in tropical landraces (HT_CAM_~0.2 and HT_CAR_~0.05) (Fig. 3 D, E, F, G, H). The outlier loci displaying the highest FST values within this region were located up to 10 kbp upstream of the *Su1* gene which is involved in the starch pathway.

Outlier FST analysis between pairs of geographic groups identified 214 and 41 regions displaying high and low differentiation, respectively (Fig. S9). BAYESCAN analysis identified 363 SNPs under selection between pair of geographic groups, including 167 new SNPs that were not previously identified between all five geographic groups (Table S8). The new highly differentiated regions identified by BAYESCAN were mostly specific to a single pair of geographic groups (Fig. S9, Fig. S10). Putative functions could be assigned to 272 of the 536 (50.7%) outlier loci identified by BAYESCAN analysis of all five and pairs of geographic groups. These included known genes involved in adaptation to abiotic stress, flowering time or human uses (Table S8 and S9).

### Genome-wide comparison of diversity between landraces and inbred lines

The panel of CK lines contained more monomorphic SNPs than landraces (263 *vs* 25) but still captured 99% of the alleles present within the landrace panel. HT was slightly higher in inbred lines than in landraces (0.353 *vs* 0.338). Allelic frequency of loci and HT values in inbred lines and landraces were strongly correlated (E=0.89 and r_2_=0.80, respectively, Fig. S11). Overall genetic differentiation between landraces and inbred lines was limited (0.010 ± 0.066). Some regions were more diverse in landraces than in inbred lines, notably the peri-centromeric region of chromosomes 3 and 7, while the opposite was found in centromeric regions of chromosomes 1, 3, 4, 5 and 6 (Fig. 3B).

Comparison of landraces and inbred lines using the outlier FST approach identified 128 highly differentiated genomic regions (FST> 0.04) and 32 regions with an excess of similarity (FST<4.21e-05). While highly differentiated regions were mainly located on chromosomes 3, 4, 8, 9 and 10, weakly differentiated regions were mainly located on chromosomes 3, 5 and 9 (Fig. 3K). BAYESCAN analysis of landraces *vs* inbred lines identified 61 loci (0.3%) that were significantly more differentiated than expected under the drift model (Fig. 3K, Table S10).

### Relationship between inbred lines and landrace populations: genetic distances and supervised analysis

The average MRD between landraces and CK lines was 0.499 (±0.034), which is greater than between landraces (0.379 ±0.059) and less than between lines (0.590 ± 0.024). The distribution of MRD genetic distances between a given landrace and CK lines (MRD_LI_) is displayed as a series of boxplots (Fig. 4A) listed in ascending order of landrace expected heterozygosity (Hs) (Fig. 4B). Landraces with a low genetic diversity generally showed a higher median and a wider range for MRD_LI_, with some notable exceptions (*e.g*. Chi5, Per10, Par2, Par1, Bra4, Ecu17, Vir4 and Svt1 in Fig. 4). Accordingly, the median MRD_LI_ and the within landrace genetic diversity Hs were strongly negatively correlated (r = −0.978, t= - 61.314, p-value < 2.2e-16) and displayed a linear relationship (Fig. S12). Considering a similar level of genetic diversity, some landraces were closely related to certain inbred lines, whereas other landraces were not (Fig. 4A and Fig. S12).

**Fig. 4:**
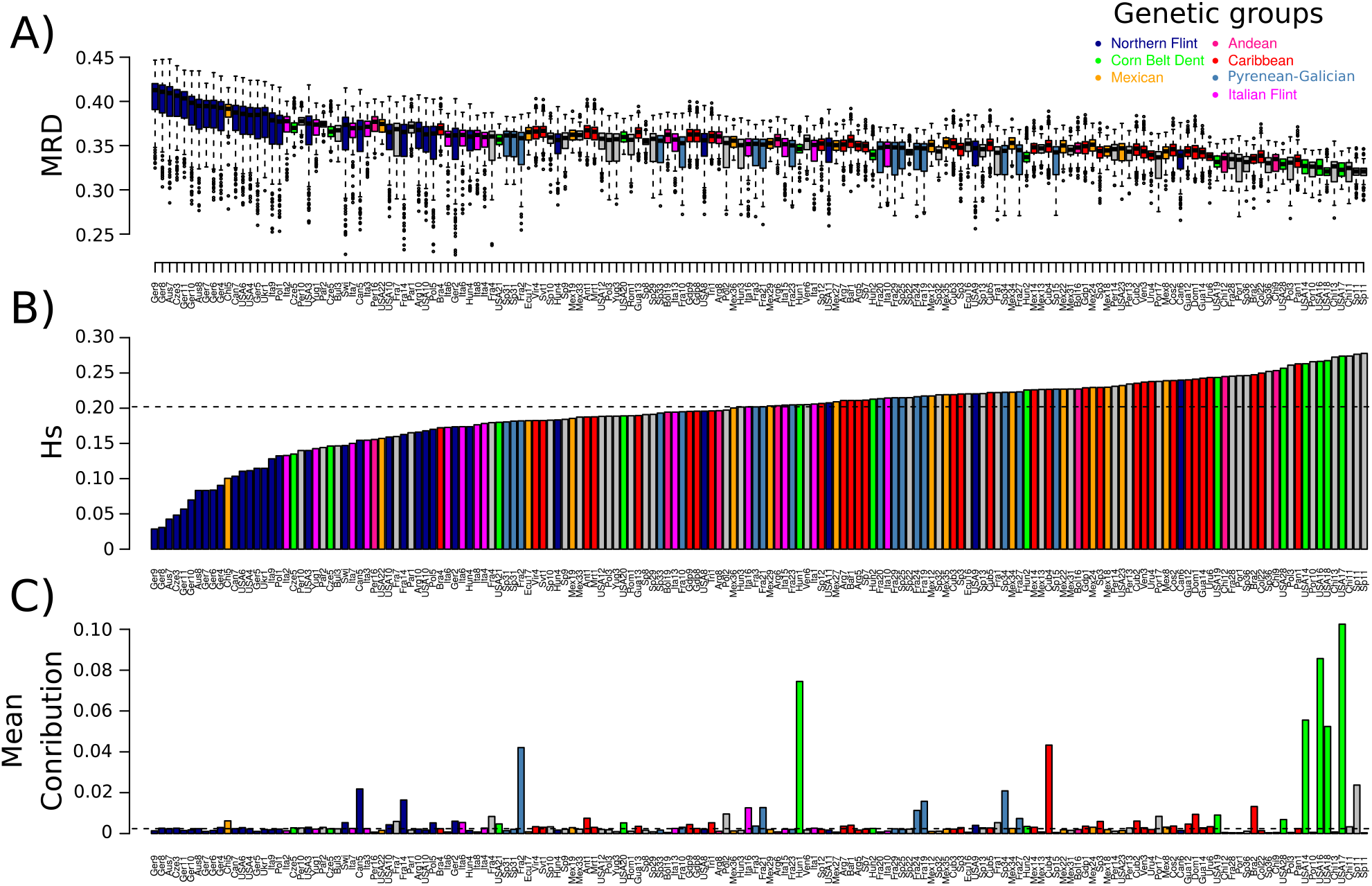
Contribution of landraces to the panel of CK lines in relation to their genetic diversity. A) Box plot representation of pairwise modified Roger’s distances (MRD) between individual landraces and inbred CK lines. Each box represents the interquartile range, the line within each box represents the median value and the error bars encompass 95% of values for each landrace. Circles represent outliers. B) Within population genetic diversity (Hs) C) Average contribution of the 166 landraces to the panel of CK lines estimated by supervised analysis with ADMIXTURE. Landraces are ranked in ascending order of Hs in the three figures. Boxplot and barplots are colored based on the assignment of landraces to the seven genetic groups identified by ADMIXTURE (see bottom right for colors).

In order to identify the source material of modern varieties, and *a contrario* the landraces that did not contribute much to these varieties, we quantitatively assigned 442 inbred lines to 166 landraces using a supervised analysis (Table S11). The 234 first cycle inbred lines (*i.e*. directly derived from a single landrace) were assigned to a total of 60 landraces. Among these landraces, 47 had at least one inbred line assigned with a probability >60%. For first cycle inbred lines of known pedigree and whose ancestral landrace is included in our study (a total of 121 lines and 50 landraces), we noted a very good match between pedigree and main assignment (71.9% of cases). Among these 121 lines, DH-SSD lines, which were derived recently from landraces, were more frequently assigned to their population of origin than lines from the diversity panel (77.6% vs 58.3%, p-value=0.04). For the 208 inbred lines from more advanced breeding cycles, we identified a total of 66 landraces as the main assignment of at least one inbred line. Among these, temperate inbred lines were frequently assigned to Reid’s Yellow Dent and Lancaster Surecrop. Chandelle (one of the few tropical landraces in our study) was identified as the most likely source for many tropical lines.

A few landraces contributed strongly to the whole diversity panel, with the 10 first landraces cumulating half of the total contributions (Fig. 4C, Fig. S13A). 80% of lines were assigned to these 10 landraces with a > 60% probability (Fig. S13B). Interestingly, the mean contribution of landraces differed strongly between first cycle lines and more advanced lines with a strong decrease (>1%) for 15 landraces and a strong increase (>1%) for 8 landraces (Fig. S13C).

We tested whether the mean contribution of landraces and the MRDLI distance “normalized” by within landraces genetic diversity could be used as a criterion to identify untapped sources of genetic diversity that could enrich the CK line panel. First, we selected 66 DH-SSD lines that were correctly assigned to 33 landraces from the landrace panel. We then classified these 33 landraces according to: (i) their average contribution to CK lines (Fig. 5A) and (ii) the normalized MRD distance from their closest lines (Fig. 5C). For each class, we estimated with 979 haplotype markers the average number of new haplotypes discovered in the 66 DH-SSD lines compared to those existing in the CK lines. We discovered 66 new haplotypes in the DH-SSD lines compared to 4,355 different haplotypes in the CK lines. The number of new haplotypes discovered in DH-SSD lines ranged from 0 (Bul3) to 11 (Arg8). The average number of new haplotypes was significantly higher for lines derived from landraces with a low contribution than those with a high contribution (p-value = 0.008, Fig. 5B). It was also higher for landraces that were not close to any of the CK lines than for those that were close to certain lines (p-value = 0.0004, Fig. 5D).

**Fig. 5:**
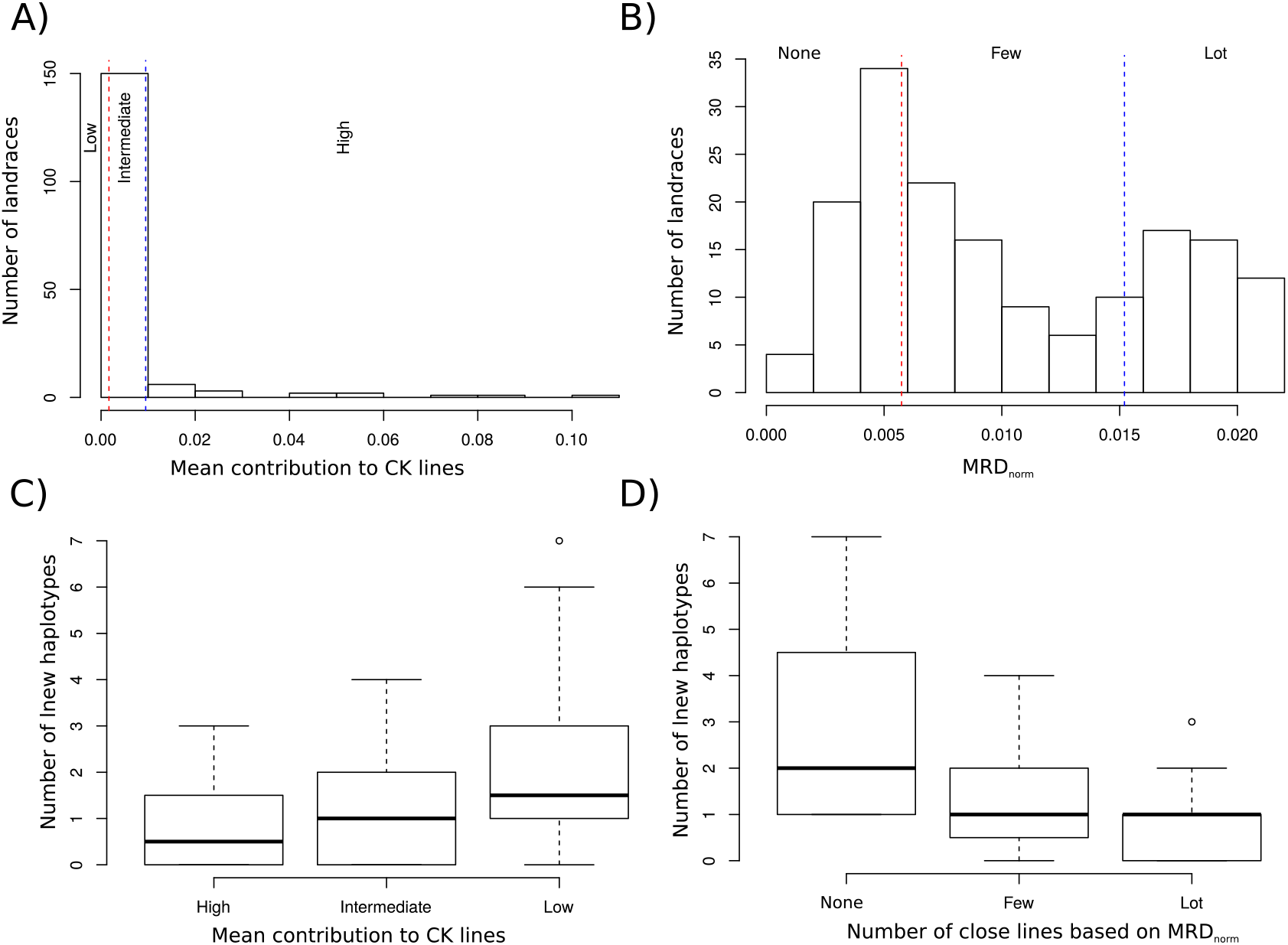
Allelic enrichment of CK lines by new DH-SSD lines derived from landraces according their contribution and their genetic distance to CK lines. Allelic enrichment was estimated by the number of new haplotypes discovered in the 66 new DH-SSD lines derived from 33 landraces, compared to the 327 CK lines (C, D) that are classified in 3 classes according to the distribution of A) the average contribution to CK line panel using supervised analysis and B) the normalized MRD (MRD_norm_) of the 10% closest CK lines with each landrace. Red and lue vertical dotted lines delineate the limits of three landrace classes displaying A) low, intermediate and high contribution; B) the presence of none, few and many closely related lines based on MRD_norm_.

## DISCUSSION

### Patterns of genetic diversity and population structure within landraces

The total expected heterozygosity observed in our study based on SNPs (0.338) was lower than the values reported previously for landraces of comparable origin that were analyzed with SSR markers (0.58 in (26), 0.63 in (27), 0.62 in (28)) but comparable to those observed with SNPs in diversity inbred line panels (42, 43). These differences can be primarily explained by the fact that SNP markers are typically bi-allelic, whereas SSR markers are multi-allelic, which has the potential to increase gene diversity (43, 52). Trends in the partition of genetic diversity within and between landraces, and within and between geographic groups were similar to previous findings. The diversity of individual landraces represented on average 57% of the total genetic diversity, which was slightly lower than for RFLP markers (~66% in (23, 26)). This difference may be due to the counter-selection of SNP markers with low MAF during the design of 50K Illumina array (37), which may increase total diversity more than within diversity (53, 54). On the other hand, genetic structure analyses based on SNPs and 17 SSRs were highly congruent, which indicates that the ascertainment bias of prefixed PZE SNPs from the 50K Illumina chip used to study landraces is negligible (43, 51).

Each geographic group contained most of the overall landrace genetic diversity, ranging from 89% (CAR) to 97% (CAM). Central American and Mexican landraces displayed the highest diversity, which is consistent with their proximity to the center of maize domestication (13, 20). This confirms that genetic diversity was lost during the spread of maize away from its domestication center due to successive bottlenecks related to climatic adaptation and isolation by distance (7, 21, 22, 29, 55). This loss of genetic diversity is consistent with the scenario of maize diffusion with (i) less genetic diversity in European than in North and South American landraces, and (ii) more diversity in South America than in North America, where maize was introduced more recently (21, 23, 29, 35). Our results nevertheless confirm that the bottleneck during the introduction of maize in Europe was certainly limited, as also shown by Brandebourg *et al*., (29) with whole genome sequencing of 67 inbred lines from Europe and America. Some northern European landraces originating from Germany and Austria have extremely low genetic diversity (Hs <0.10), with more than 70% of loci being fixed, suggesting a strong bottleneck. The fact that some of these landraces have been cultivated mostly in gardens may have decreased their effective population size (26). The genetic load could have been more or less purged depending on the severity and the duration of the bottleneck. This could explain the strong variation in success rate observed for deriving inbred lines from European Flint landraces by haplodiploidization (56, 57).

Genetic distance, Ward hierarchical clustering (Fig. 1B), principal component (Fig. 1A) and population structure (Fig. 2) analyses showed major trends in population structure. We confirmed the central position of Mexican and Caribbean landraces and a clear differentiation between North and South American landraces (Fig. 1 and 2). This is consistent with the domestication of maize in Mexico followed by southwards and northwards dispersion (22, 55). The similarity between landraces from southern Spain and the Caribbean confirms the historical data on the introduction of maize in the south of Spain by Columbus in 1493 after his first trip to the Caribbean (Fig. 1B, cluster d). Strong similarities between groups of northeastern American and northeastern European landraces (mostly from Germany, Poland and Austria) (Fig. 1B, cluster g) also supports an independent introduction of North American material that was pre-adapted to the northern European climate (21, 22, 26, 28, 29, 58). Some landraces from northern Spain and southwestern France, located along the Pyrenean Mountains, were admixed either with Caribbean or Northern Flint. This result supports the hypothesis that new Pyrenean-Galicia Flint groups originated from hybridization between Caribbean and Northern Flint material that were introduced in southern Spain and northern Europe, respectively. (27, 29, 59). Interestingly, some southwestern Spanish landraces have elevated admixture with Italian Flint groups and are closely related to Italian landraces on the NJ tree (Fig. S5), while northern Spanish landraces (latitude >42°N) do not. These results support the hypothesis that Italian landraces are probably derived from an ancestor from southern Spain (29, 60). Our results also highlighted a new putative hybridization event in Central Eastern Europe. Central Eastern European landraces were close to Italian Flint landraces on the Ward cluster tree and one northern Italian Flint landrace (Nostrano Quarantino) was admixed with Italian Flint (~30-40%) and Northern Flint (~30-50%). This suggests that Italian Flint landraces certainly spread in Central Eastern Europe, where they intermated with Northern Flint landraces.

Differentiation of landraces was greater in Europe than in Central America and the Caribbean, indicating that gene flow is lower in the latter two. Genetic and geographic distances were significantly correlated in NAM, EUR and CAM but not in SAM and CAR (Fig. S6), suggesting that isolation by distance played a role in shaping the genetic structure of maize landraces in these regions, albeit to a variable degree. In the case of CAM, the effect of isolation by distance is partially blurred by variation in altitude producing major gradients in environmental conditions (temperature, rainfall) (7, 30, 61). Indeed, Mexican landraces clustered according to both altitude and distance (Fig. 1B, Table S1) suggesting environmental adaptation (7, 30).

### Genomic pattern of nucleotide variation in landraces

FST outlier and BAYESCAN analyses identified 13 genomic regions that showed high levels of differentiation between geographic groups and/or landraces (Table S5). The four highly differentiated genomic regions between landraces displayed contrasted patterns of allelic frequencies between geographic groups (Table 2, Table S5), suggesting different types of selection. The Sp10 region was found to be highly differentiated between landraces but not between the five geographical groups. It suggests that there was strong selection in some specific geographic areas but not across all geographic groups. This region contains genes associated with tolerance to high temperature and evaporative demand (62). The second genomic region (Sg4-Sp6: 7.8 Mbp – 9.3 Mbp on chromosome 4) was nearly fixed in temperate landraces (NAM, EUR) whereas it showed intermediate frequencies in CAM, suggesting a strong directional selection effect during the spread from Mexico to North America. This results is in agreement with Romero-Navaro et al. (55), who identified 5 SNPs in this region with allelic frequencies varying significantly with latitude in American landraces, and Brandeburg et al., (29), who identified two highly differentiated regions between Corn Belt Dent and Tropical first cycle lines. By contrast, the third genomic region (Sp5-Sg7; 40-41.9 Mbp on chromosome 4) displayed higher genetic diversity in temperate landraces (NAM, EUR) than in tropical landraces (CAM, CAR) suggesting strong diversifying selection in EU and NAM. This region included the *Su1* gene, which is involved in the starch pathway and is known to be under strong selective pressure (63–66). Romero-Navaro et al., (55) also found an association between allelic frequency variation at the *Su1* locus and both latitude and longitude. Futhermore, Brandeburg et al., (29) identified a strong selective sweep between Corn Belt Dent/Tropical and Northern Flint first cycle lines in the *Su1* gene. The fourth region (Sg2-Sp3; 84-85 Mbp on chromosome 3) showed a continuous gradient of allelic frequencies between tropical and temperate landraces suggesting strong directional selection for adaptation either to temperate or tropical climates. In agreement with this finding, Romero-Navaro et al., (55) identified in this region 22 and 4 SNPs with allelic frequencies varying significantly with altitude and latitude, respectively. This region also carries a large 6 Mbp inversion that is putatively involved in flowering time variation (55).

BAYESCAN analysis between geographic groups identified several regions that were not identified by outlier FST analysis (Table S8 and S9). Notably, we identified several loci under strong selection that were close to genes known to be involved in flowering time variation: (i) PZE-108070380 on chromosome 8 (123.5 Mbp) localized 5 kbp upstream of *Zcn8* (42, 67, 68); (ii) PZE-109070904 on chromosome 9 (115.7 Mbp) in *ZmCCT9* (69); (iii) two loci on chromosome 3 (PZE-103098664 (158.9 Mbp) and PZE-103098863 (159.17 Mbp) close to *Vgt3*, a major loci that is strongly associated with flowering time variation in temperate maize (62, 70). We also identified several genes/genomic regions that are putatively involved in adaptation to abiotic stress: (i) PZE-102108435 on chromosome 10 that is 10 kbp upstream of *ZmASR2* which is involved in abscisic stress ripening (71); (ii) PZE-104128228 on chromosome 4 in the *nactf125* gene (within Sg6 in table S5), PZE-102051809 in the *nactf36* gene (chromosome 1) and PZE-107058109 in the *nactf14* gene (chromosome 7), all of which belong to the NAC protein family, which encodes plant transcription factors involved in biotic and abiotic stress responses (72); (iii) two diaglycerol kinases (*dgk2* and *dgk3*) that exhibit differential expression patterns in response to abiotic stress including cold, salinity and drought and are upregulated in cold conditions (73). Finally, we identified several genomic regions carrying genes involved in the hormonal systems regulating growth, cell division and proliferation such as giberellin2-oxydase9 (*ZmGA2ox9*, GRMZM2G152354), phytosulfakine (GRMZM2G031317) or in the starch pathway (*Su1, waxy1, dull endosperm1*).

The detection of genomic regions and loci under selection have therefore allowed the identification of genes that underlie the adaption of maize to diverse agro-climatic conditions and/or human uses during the spread of landraces from America (7, 22, 23, 29, 55, 74). These genomic regions could be useful for mining new alleles from landraces, retrieving some of the genetic diversity that was lost by genetic drag linked to genes close to those under selection (7, 41, 74), or creating new genetic diversity by targeted mutation (7).

### Identification of promising landraces to enlarge the modern genetic pool

Intensive selection to enhance agronomic performance can considerably reduce genetic diversity in crops (1). However, we found little difference in genetic diversity between landraces and inbred lines, which is consistent with the low genetic differentiation we observed between landraces and inbred lines. This suggests that the genomic diversity (inferred from SNPs) present in landraces was retained in our panel of CK lines and that selection during maize improvement has not altered allele diversity over a very broad geographic scale. This observation is similar to findings in soybean (75) and wheat (76), which also showed a minor effect of crop improvement on diversity, suggesting that landraces have been and still are extensively used in the development of modern inbred lines in these crops. It is important to note however that our line panel included many old lines that have made only a limited contribution, if any, to commercial F1 hybrids or recent breeding pools. Our panel therefore certainly overestimates the genetic diversity present in the germplasm of modern breeding inbred lines (57).

Several factors could be responsible for the low genetic erosion accompanying the transition from landraces to inbred lines. A first hypothesis is that selection during modern maize breeding targeted only a small number of genes (77) and therefore affected genetic diversity and allelic frequency only in the genomic regions flanking the genes under selection. Another hypothesis is that, even if only a limited number of landraces were used as parents of first cycle lines, i.e. the initial modern inbred line breeding pools, selection of genetically diverse and complementary heterotic groups may have mitigated the loss of diversity (78). Furthermore, SNPs from 50K arrays were previously identified in 27 lines (79). These SNPs may not reflect well the total genetic diversity of landraces, as certain specific landrace haplotypes may not have been transmitted to first cycle lines due to their deleterious effect at the homozygous state (inbreeding depression) or gamete sampling (drift) (57).

Despite the limited differences in overall diversity between landraces and inbred lines, two different approaches highlighted that the majority of landraces had made a limited contribution to recent breeding. We identified a number of landraces with a high median Hs value and a small MRD_LI_ distance range reflecting a lack of similarity similarity to any inbred line. These landraces probably did not contribute to the modern maize germplasm. Indeed, supervised analysis showed that inbred lines from our diversity panel could be traced back to a few landraces and that the first 10 landraces cumulated half of the total contribution to the diversity panel. Most of these landraces (Reid’s Yellow Dent, Lancaster Surecrop and Krug Yellow Dent for the dent genetic group, Lacaune and Gaspe Flint for the flint genetic group and Chandelle for Tropical lines) were previously identified as the source of the modern maize breeding germplasm (12, 13, 55). Interestingly, we observed a large increase or decrease in the contribution of landraces between first cycle lines and more advanced lines (Fig. S13C). This can be explained by the fact that some lines were extensively used to derive more advanced lines whereas others were not (12, 14, 15). Interestingly, DH-SSD lines that were recently derived from landraces were assigned more frequently (and with higher probability) to their population of origin than older lines that were maintained for a long time in gene banks. This suggests that some landraces could have evolved since contributing to inbred lines from the diversity panel or that the pedigree of these lines was erroneous. Our results suggest that we could use supervised analyses to curate the landrace collection and the pedigree of first cycle lines.

In order to identify landraces that differ the most from inbred lines, we developed an indicator of genetic distance from inbred lines which was normalized by their genetic diversity (Fig. S12). By classifying landraces according to (i) this normalized distance and (ii) their average contribution to reference inbred lines, we were able to identify landraces that have the greatest potential to broaden the genetic diversity of these lines (Fig. 5). By combining closely located SNPs, we were able to identify novel haplotypes in the DH-SSD lines, which were absent in the CK panel, even though both alleles were present in landraces and the inbred line panel. The number of new haplotypes was significantly higher for DH-SSD lines created from landraces classified as genetically distant from the modern germplasm according to the criteria described previously, which confirms their relevance when choosing landraces for diversity enhancement. This strategy to identify untapped landraces in modern breeding germplasm can be easily extended to other plant species, other material (hybrids, private germplasm), and other technologies (sequencing). Additionally, this strategy can be focused on some genomic region to identify new alleles of interest. Our strategy opens an avenue to identify valuable landraces and genomic regions for prebreeding.

## MATERIALS AND METHODS

### Plant material

#### Landraces

A total of 156 different landrace populations (Table S1) were sampled from a panel of 413 landraces (Supplementary Information 1). These 156 landraces captured a large proportion of European and American diversity and have been analyzed in previous studies using RFLP (25, 31–34) and SSR markers (23, 27, 28). Each population was represented by either one or two sets of 15 individual plants (for 146 and 10 populations, respectively), pooled equally as described in Reif *et al*. (48) and Dubreuil *et al*. (28). The 166 DNA samples corresponding to the 156 landrace accessions were classified into five geographic groups (Table S1): Europe (EUR), North America (NAM), Central America and Mexico (CAM), the Caribbean (CAR) and South America (SAM).

#### Inbred lines

We analyzed 234 inbred lines that were derived directly by single seed descent or by haplodiploidization of landraces, referred to as “first cycle lines”, and 208 lines that were derived from a more advanced cycle of breeding, referred to as “advanced lines” (Table S11). These 442 lines were partitioned into three sets (the “Panel” column in Table S11):

1. “CK lines”: a panel of 120 first cycle and 207 advanced lines (327 lines in total) representing American and European diversity (27, 42) including some key founders of modern breeding programs (*e.g*. F2, B73, C103).
2. “Parent Controlled Pools”: a set of 12 lines used to build 4 series of 8 controlled DNA pools (see below).
3. “DH-SSD lines”: a set of 45 single seed descent (SSD) and 58 double haploid (DH) lines derived recently from 48 landraces (first cycle lines).

#### Controlled DNA Pools

To prepare the controlled DNA pools, two sets of three inbred lines were considered: EP1 – F2 – LO3 (European Flint inbred lines) and NYS302− EA1433 – M37W (Tropical inbred lines). For each set of parental lines, nine controlled pools were prepared by varying the proportion of each line in the mix, quantified by the number of leaf disks of equal size as *per* Dubreuil et al., (32). The genotype of each line and the proportion of parental lines in each pool were used to estimate allelic frequencies in the nine pools, and subsequently to calibrate the model for predicting allelic frequency (see (51) for more detail).

#### Genotyping and prediction of allelic frequencies in DNA pools

We used the 50K Illumina Infinium HD array (37) to genotype (i) landraces, (ii) controlled DNA pools, (iii) the DH-SSD inbred lines and (iv) the parental lines of the controlled DNA pools(Table S1 and S11). For CK lines, we used the 50K genotyping data from Bouchet et al. (2013). 23,412 SNPs were filtered based on their suitability for diversity analysis and their quality for predicting allelic frequency in DNA pools (Supplementary Information 2).

Allelic frequency of selected SNPs in DNA pools was estimated using the two-step procedure described in (51) based on the fluorescence intensity ratio (FIR) of alleles A and B for each SNP. First, we tested whether SNPs were monomorphic or polymorphic. For SNPs that were considered to be polymorphic, we then estimated the allelic frequency of the B allele using a generalized linear model calibrated on FIR data from 1,000 SNPs from 2 series of controlled pools (see (51) for more detail and equation 2 for the model).

We also used the genotyping data from 17 SSRs from 145 and 11 landraces obtained by Camus-Kulandaivelu et al. (27) and Mir et al. (23), respectively.

### Diversity analyses

#### Estimation of genetic diversity parameters

For each landrace, each geographic group, all landraces combined and the panel of inbred lines, we determined for each locus: the mean allele number (A), the Minor Allele Frequency (MAF) and the expected heterozygosity (H) (80, 81).

Genetic differentiation (FST) was estimated between: individual landraces (FSTl), between the five landrace geographic groups (FST_g_), between 10 pairs of geographic groups (FST_EUR-NAM_, FST_EUR-CAM_, FST_EUR-CAR_, FST_EUR-SAM_, FST_NAM-CAM_, FST_NAM-CAR_, FST_NAM-SAM_, FST_CAM-CAR_, FST_CAM-SAM_, FST_CAR-SAM_) and between landraces and inbred lines (FST_i_). FST was estimated at each locus and across all loci as *per* (81, 82) (Supplementary Information 3).

#### Genome-wide diversity analysis and scans for identifying selection signatures

We used a sliding window of 1 Mbp, shifting by 500 kbp at each step along the genome, to analyze the genome-wide variation in genetic diversity and differentiation between landraces, between geographic groups, and between landraces and inbred lines. The maize genome was divided into 4,095 overlapping windows containing an average of 11.3 ± 5.2 SNPs. We computed the average value for the parameters described above for all loci in a given window. Outlier regions for H and FST were identified based on the distribution of these parameters for individual loci over the entire genome using the 5th and 95^th^ percentile (below 5% and above 95%) as thresholds (Table S4). All statistics were computed using *ad hoc* scripts in the R language v 3.0.3 (83).

Genomic scans were carried out to detect the genomic signature of selection between landraces, between the five geographic groups and between landraces and inbred lines using two approaches: (i) the detection of 1 Mbp regions that were outliers for FST, referred to as “Outlier FST analysis” (ii) the detection of loci under selection using the drift model implemented in the BAYESCAN software (84) (Supplementary Information 4).

#### Genetic structure and relationship between landraces

We estimated the genetic distance between all landraces using modified Roger’s distance (MRD) (85) based on the allelic frequencies of 23,412 prefixed PZE SNPs. MRD was then averaged within and between geographic groups (Table 1, Table S2). We analyzed the relationship between genetic and geographic distances within each geographic group by plotting MRD against geographic distances. We tested this correlation using the Mantel test (86). Geographic distances were calculated using the latitude and longitude of each sampling site using the geosphere R package v. 1.5-10 (87).

To decipher the structure of genetic diversity within our panel of landraces from 23,412 filtered SNPs, we used two approaches:

1. A distance-based approach in which MRDs between the 166 landraces were used to perform (i) a principal coordinate analysis (PCoA) (88), (ii) hierarchical clustering using either Ward or Neighbor-Joining algorithms implemented in the “hc” and “bionj” functions of the “ape” R package v 5.0 (89), respectively.
2. A Bayesian multi-locus approach, implemented in the ADMIXTURE software, to assign probabilistically each landrace to K ancestral populations assumed to be in Hardy-Weinberg Equilibrium (90). Different methods were used to identify the most appropriate number of ancestral populations (K): Cross-validation error or difference between successive cross-validations (90) and Evanno’s graphical methods (91). Since ADMIXTURE requires multi-locus genotypes of individual plants, we simulated the genotype of five individuals for each population for a subset of 2,500 independent SNPs to avoid artifacts of linkage disequilibrium (Supplementary Information 5).

### Contribution of populations to inbred lines using supervised analysis and modified Roger’s distance

To analyze the contribution of landraces to the modern breeding germplasm, we used two different approaches:

1. A distance-based approach in which we estimated the modified Roger’s distance between each landrace and the 327 CK lines (MDRLI) in order to determine whether they are related or not.
2. A Bayesian supervised approach implemented in ADMIXTURE in which the 442 inbred lines were assigned probabilistically to the 166 landrace populations in order to identify the most likely source population of each inbred line (Table S11). For each landrace, we estimated (i) its average contribution to CK lines by averaging the assignment probability over 327 lines and (ii) the number of inbred lines mainly assigned to this landrace, with an assignment probability > 60%. We also analyzed the evolution of the contribution of landraces across breeding cycles by comparing contributions to (i) first cycle lines and (ii) advanced lines from the CK line panel. To check the accuracy of the assignment method, we estimated the percentage of first cycle lines that were correctly assigned to their parental landrace as known from their pedigree and analyzed in our study (121 of the 234 first cycle lines, known to be derived from 50 landraces). We tested if this percentage was different between CK lines and DH-SSD lines using a Kruskal-Wallis chi-squared test. To represent each landrace, we used the same five simulated individuals as in the structure analysis.

Identification of landraces that could enrich the modern breeding germplasm We assessed whether the mean contribution of landraces and their MRD_LI_ distribution parameters could be used as criteria to identify landraces that could enrich the modern breeding germplasm. To this end, allelic diversity was estimated in the two inbred panels (DH-SSD and CK lines) for 979 haplotypes. These haplotype markers were defined by genotyping triplets of adjacent SNPs from 50K arrays that were less than 2 kbp apart. We estimated the average number of new haplotypes discovered in the DH-SSD lines compared to those in the 327 CK lines. To avoid noise due to seedlot error during DH-SSD line production, we selected 66 DH-SSD lines that were correctly assigned to 33 landraces analyzed from this study.

To analyze the effect of mean contribution, we classified these 33 landraces into three classes: low, intermediate, and high contribution based on the 30th and 90^th^ percentile of the distribution of mean landrace contribution to CK lines.

To analyze the usefulness of MRDLI, we took into account the negative correlation between MRD_LI_ and within-gene diversity of landraces (Hs), which could strongly bias against landraces with the lowest within diversity. For each landrace, we defined a “normalized” MRD distance (MRD_norm_) based on the absolute difference between (i) the median MRD_LI_ between a landrace and lines of CK panel (MRD_med_) and (ii) the MRD_LI_ from the closest lines (MRD_q_) defined by the 5th (MRD05) and 10^th^ (MRD10) percentile of MRD_LI_, corresponding to the 5 and 10% closest lines. In order to correct the bias due to Hs, we used the linear regression coefficient “a” between MRD_med_ and Hs. We defined MRD_norm_ as the orthogonal deviation of MRD_q_ (with q = 5% or 10% for MRD_05_ and MRD_10_, respectively) from the linear regression:

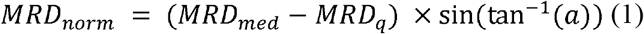

We used MRD_norm_ based on MRD_10_ to categorize the 33 landraces into three classes based on the percentile distribution of MRD_norm_. Landraces with MRD_norm_ below 30%, between 30% and 70% quantile and above 70% were considered to have none, few or many derived lines, respectively.

Finally, we performed a variance analysis to test the effect of mean contribution and MRD_norm_ on the number of new haplotypes discovered in the DH-SSD lines.

## Acknowledgements

This study was funded by the” Association pour l’étude et l’amélioration du maïs” (PROmais) within the “Diversity Zea” project and the French National Research Agencies with thein “Investissement d’Avenir Amaizing” project, (ANR-10-BTBR-01). We greatly acknowledge the French Maize Biological Ressource Center, PROmais, and the INRAE experimental units of St Martin de Hinx and Mauguio for collecting and maintaining the collection of landraces and inbred lines. We greatly acknowledge colleagues who initially collected these landraces and André Gallais for initiating these research programs. We also greatly acknowledge Pierre Dubreuil, Letizia Camus-Kulandaivelu, Cecile Rebourg, Céline Mir, Domenica Maniccaci who conducted previous studies on these landraces using the DNA pooling approach with SSR and RFLP markers. The Infinium genotyping work was supported by CEA-CNG. We thank Anne Boland and Marie-Thérèse Bihoreau and their staff. We acknowledge the EPGV group, Dominique Brunel, Aurélie Bérard and Aurélie Chauveau for discussion and management of Illumina genotyping.

## Author contributions

S.D.N, A.C and B.G designed and supervised the study and selected the plant material; M.A, S.D.N, A.C, B.G drafted and corrected the manuscript; D.M, V.C and M-C.L-P extracted DNA and managed the genotyping of landraces and inbred lines; C.B, B.G and A.C collected andmaintained the collection of landraces and inbred lines; S.D.N, M.A, A.C and T.M-H developed the statistical methods for predicting allelic frequency from fluorescence data; M.A, B.G and S.D.N analyzed the genetic diversity of the landrace panel; M.A and S.D.N analyzed the selective sweep; MA, S.D.N and A.C investigated the relationship between landraces and inbred lines; S.D.N developed the normalized distance measure and performed the analysis of diversity enrichment.

## Data availability

Fluorescence Intensity Data from 166 DNA samples of landraces used for predicting allelic frequency and modified Roger’s distance matrix are available at https://doi.org/10.15454/D4JTKB. To predict allelic frequency in 166 DNA pools, we calibrated our two-step model with fluorescence intensity data of 327 inbred lines (for calibrating the fixation test) and two series of controlled pools (for calibrating logistic regression) with R script that are available at the following address: https://doi.org/10.15454/GANJ7J. Note that data will be available at the two web links below upon the publication will have been accepted in a journal.

