## Supplementary material for "Genome-wide SNP genotyping of DNA pools identifies untapped landraces and genomic regions that could enrich the maize breeding pool": Fig. S1

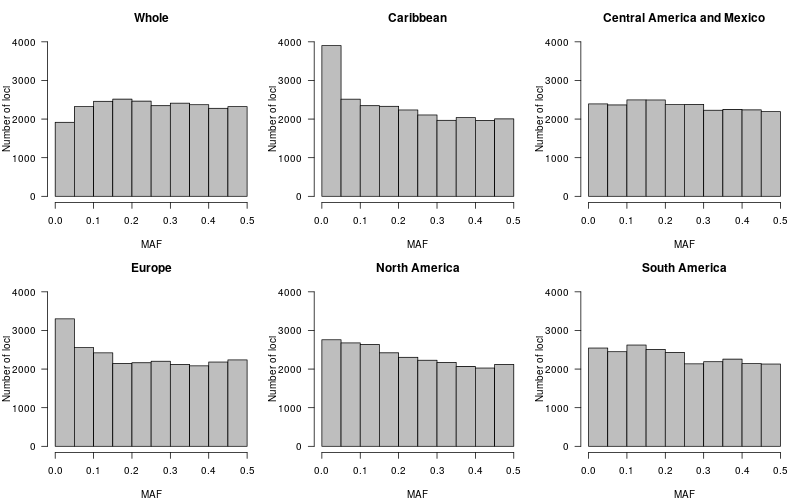
Fig. S1: Distribution of the minimum allele frequency (MAF) across the entire landrace panel (Whole) and within the five geographic groups.
