## Supplementary material for "Genome-wide SNP genotyping of DNA pools identifies untapped landraces and genomic regions that could enrich the maize breeding pool": Fig. S2

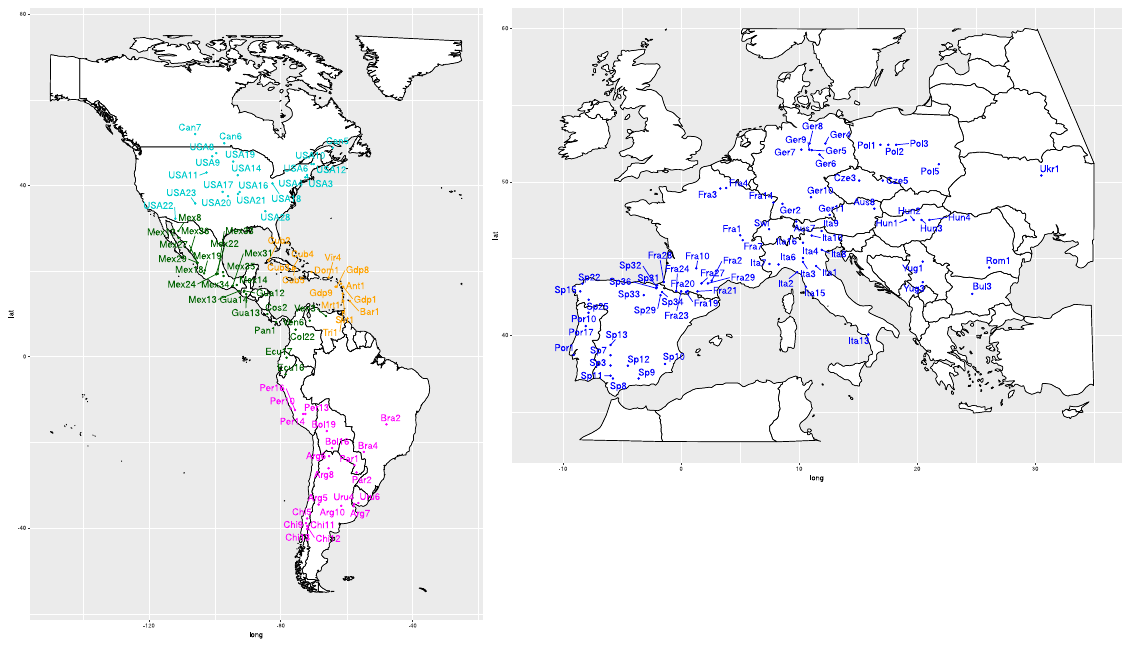

#### Fig. S2: Geographic distribution of the 156 landrace accessions. Landraces are colored according to their geographic origin: Europe (blue), North America (cyan), South America (magenta), Central America and Mexico (dark green) and the Caribbean (orange). When the exact geographic coordinates of a landrace were unknown, we used the coordinates of the city representative of the country or region where the population was collected.
