## Supplementary material for "Genome-wide SNP genotyping of DNA pools identifies untapped landraces and genomic regions that could enrich the maize breeding pool": Fig. S3

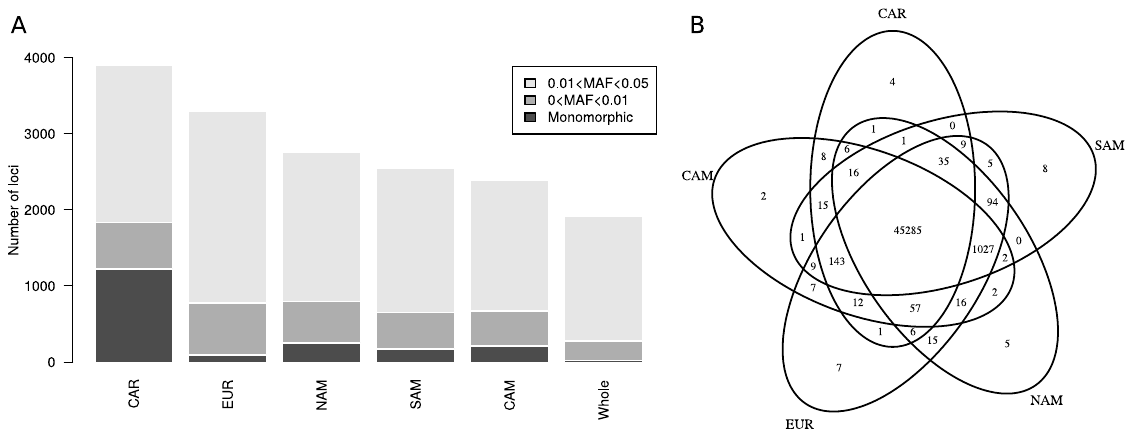

#### Fig. S3: Distribution of rare alleles in the five geographic groups of maize landraces. A) Number of rare alleles in the five geographic groups of maize landraces. For each bar, the height of the colored segment represents the number of alleles that are absent (black), with a frequency < 0.01 (dark gray) or between 0.01 and 0.05 (light gray). B) The Venn diagram illustrates the number of alleles that are exclusive to the various combinations among the five geographic groups of maize landraces.
