## Supplementary material for "Genome-wide SNP genotyping of DNA pools identifies untapped landraces and genomic regions that could enrich the maize breeding pool": Fig. S4

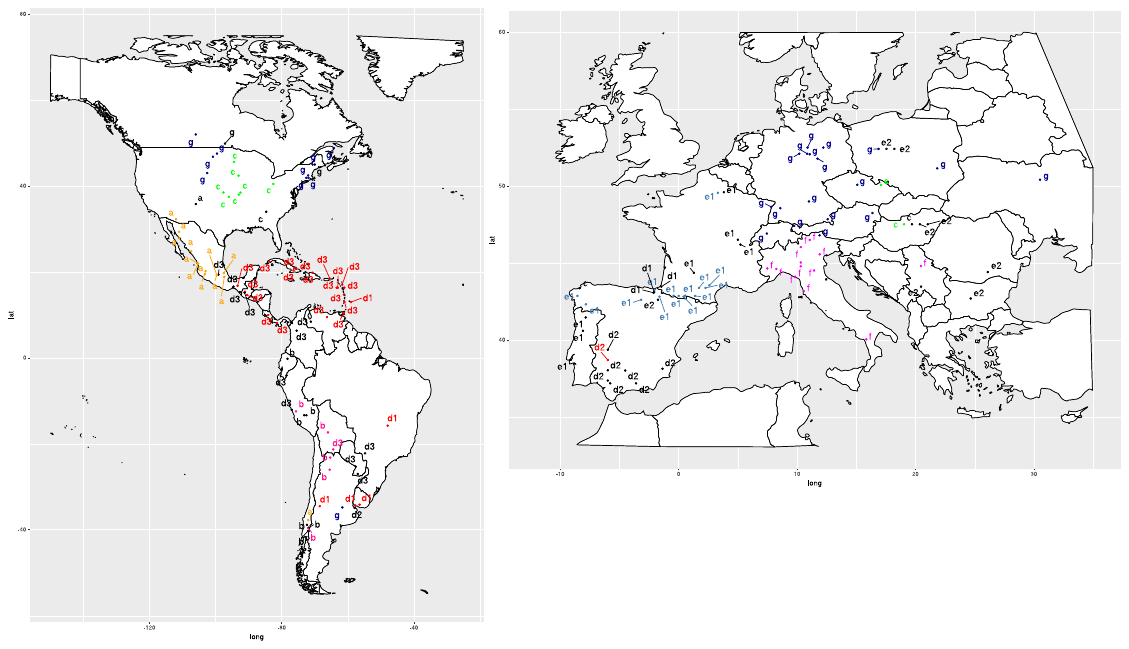

#### Fig. S4: Geographic distribution of landrace clusters obtained by hierarchical clustering based on modified Roger’s distance and Ward’s method. Landrace clusters correspond to those described in Fig. 1A and Table S1. Each landrace is colored by genetic group based on its assignment probability > 0.6: Northern Flint (dark blue), Corn Belt Dent (green), Mexican (orange), Andean (deep pink), Caribbean (red), Pyrenean-Galician (steel blue) and Italian (magenta). Admixed landraces are colored in black.
