## Supplementary material for "Genome-wide SNP genotyping of DNA pools identifies untapped landraces and genomic regions that could enrich the maize breeding pool": Fig. S5

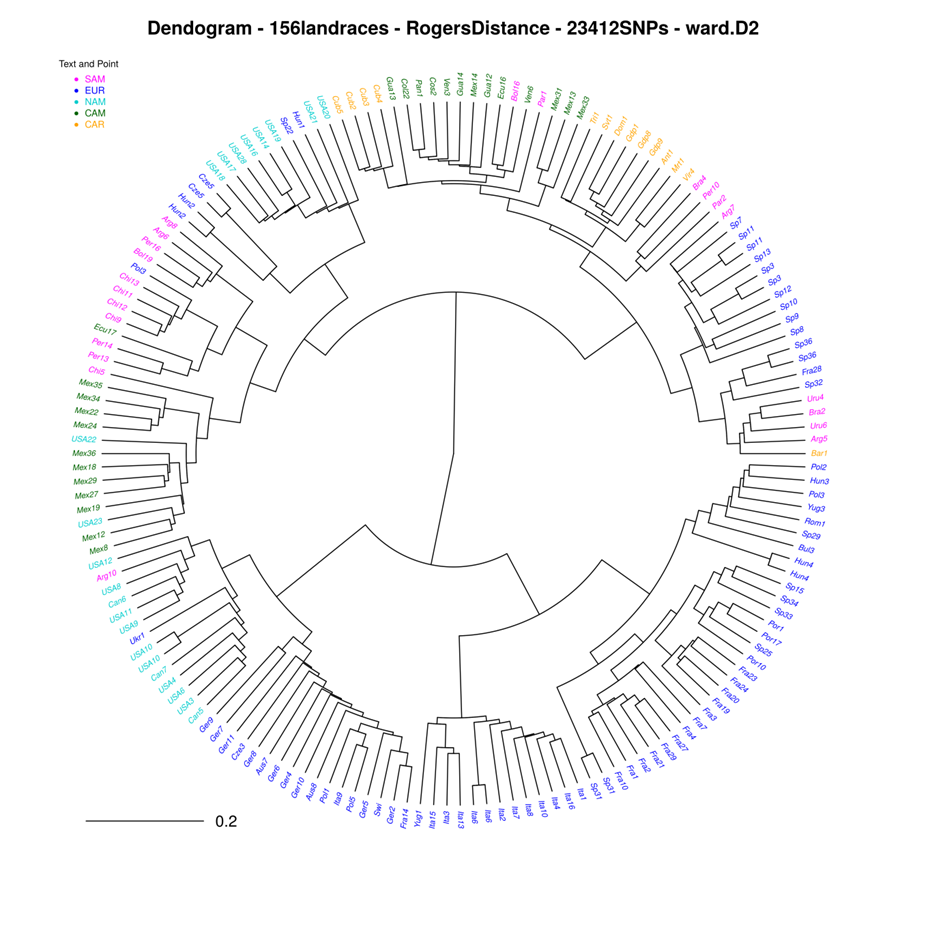

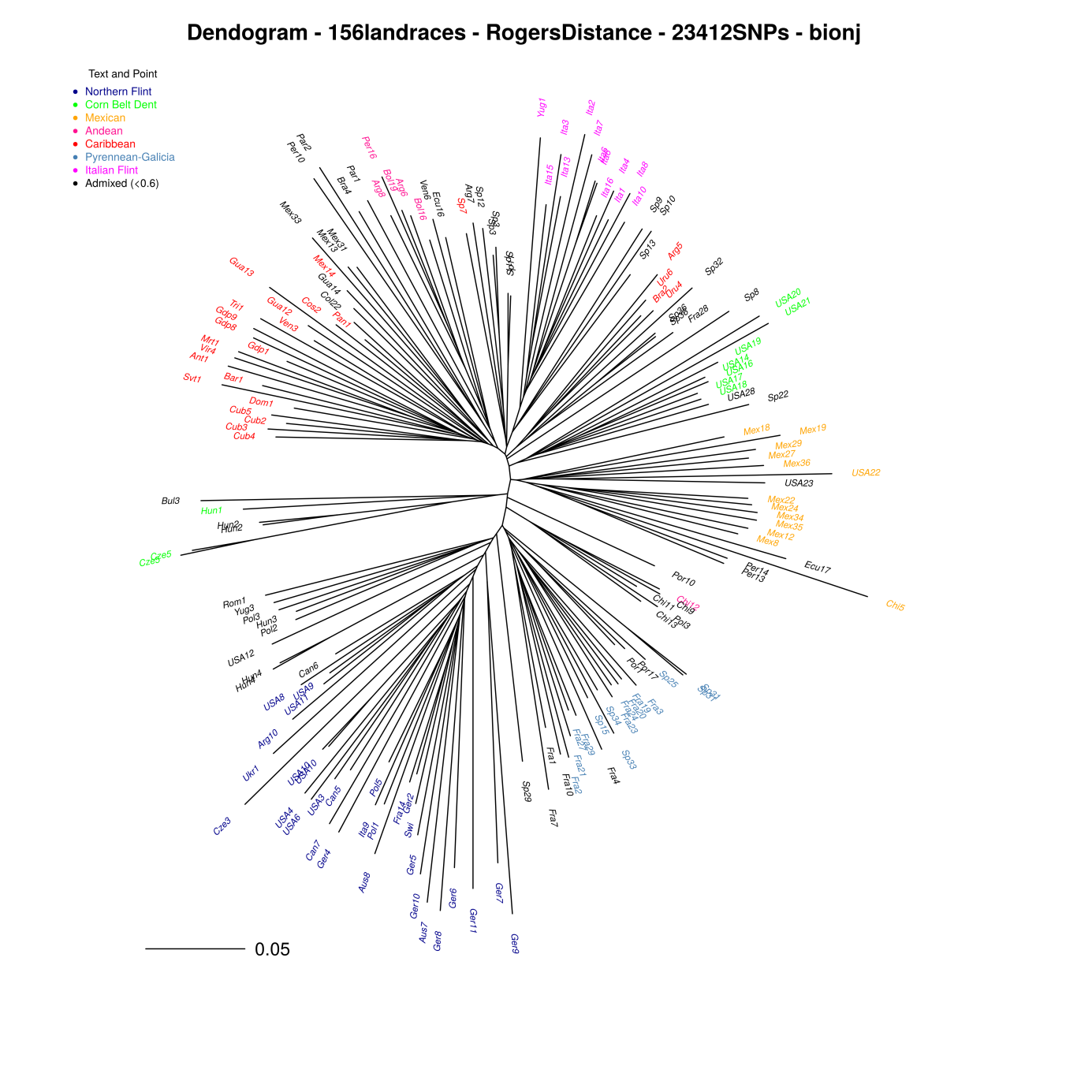

)

B)

#### Fig. S5: Dendrogram of the 166 DNA samples corresponding to 156 landrace accessions based on Ward (A) and Neighbor-Joining (B) hierarchical clustering. DNA landrace samples are colored based on their assignment to genetic groups defined at K=7 by the Admixture software for Neighbor-Joining and on their geographic origin for hierarchical clustering. Only accessions with an assignment > 0.6 are colored, whereas admixed accessions (<0.6) are indicated in black).
