## Supplementary material for "Genome-wide SNP genotyping of DNA pools identifies untapped landraces and genomic regions that could enrich the maize breeding pool": Fig. S6

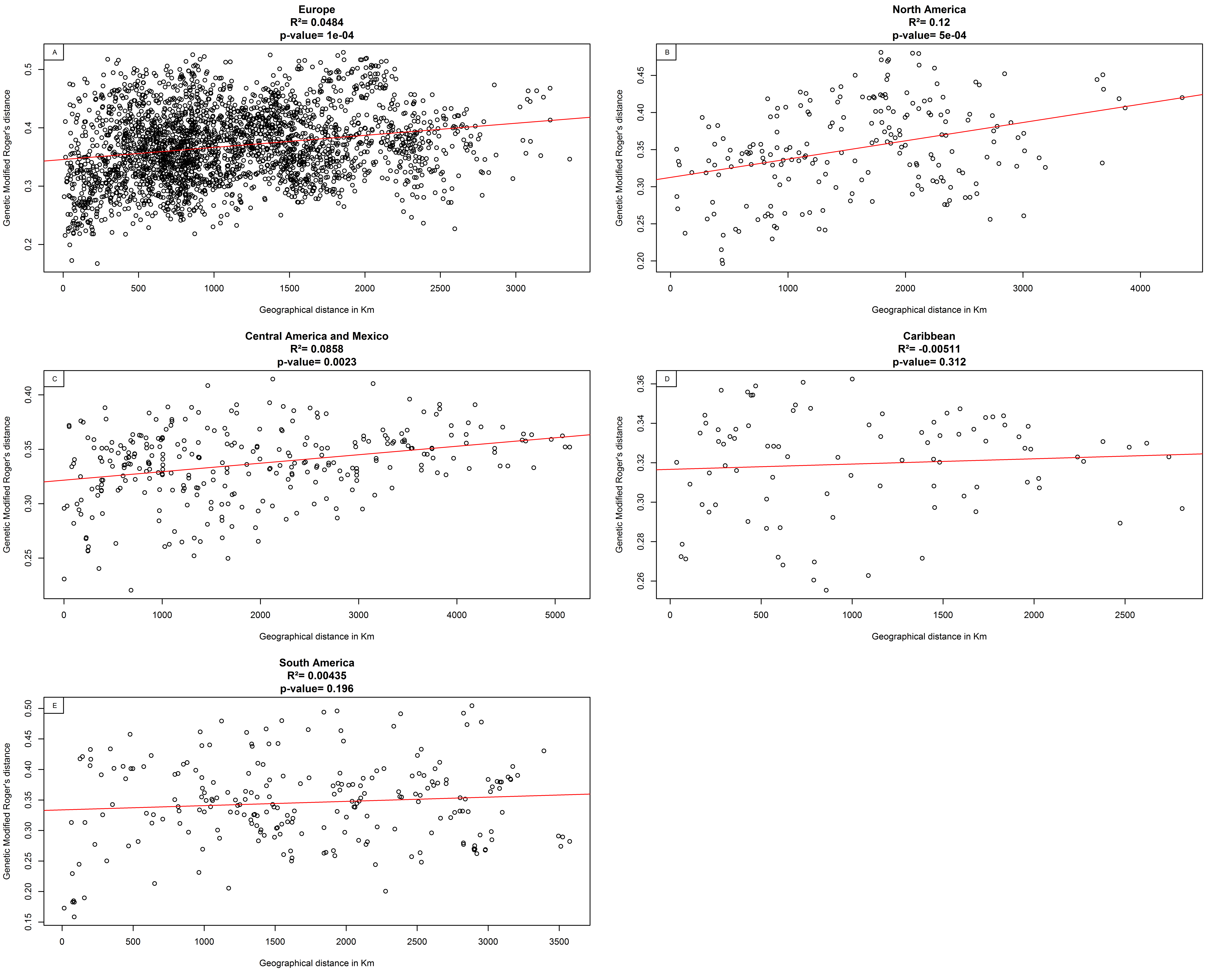

#### Fig. S6: Relationship between modified Roger’s genetic distances and geographic distances between landraces within the five geographic groups: A) Europe (EUR) B) North America (NAM), C) Central America (CAM), D) the Caribbean (CAR), F) South America (SAM). Each point represents a pair of landraces. Coefficient of determination (r^2^) and p-value of the Mantel test are indicated above each plot. Red lines represent the linear regression between MRD and geographic distance.
