## Supplementary material for "Genome-wide SNP genotyping of DNA pools identifies untapped landraces and genomic regions that could enrich the maize breeding pool": Fig. S7

#### Fig. S7: Determination of the K value by ADMIXTURE analysis performed across the landrace panel for 2,500 Panzea markers. A) Cross validation errors estimated for K values ranging from 2 to 14; B) Difference between successive cross-validation error values; C) Evanno’s graphical method with Mean L(K) over 20 runs for each K value. D) ∆K calculated as per Evanno et al. (2005).

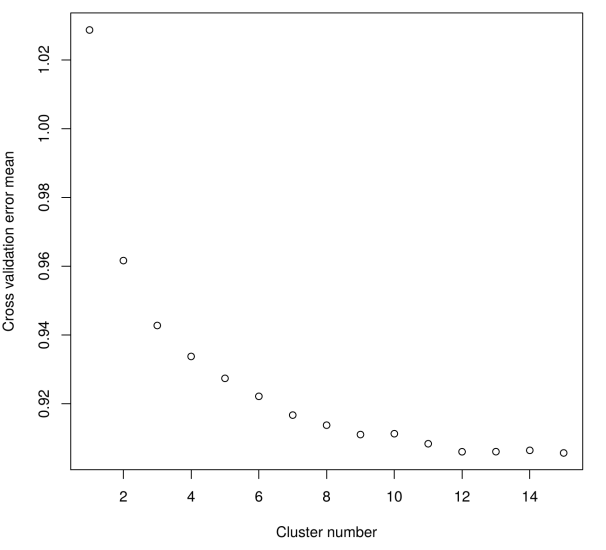

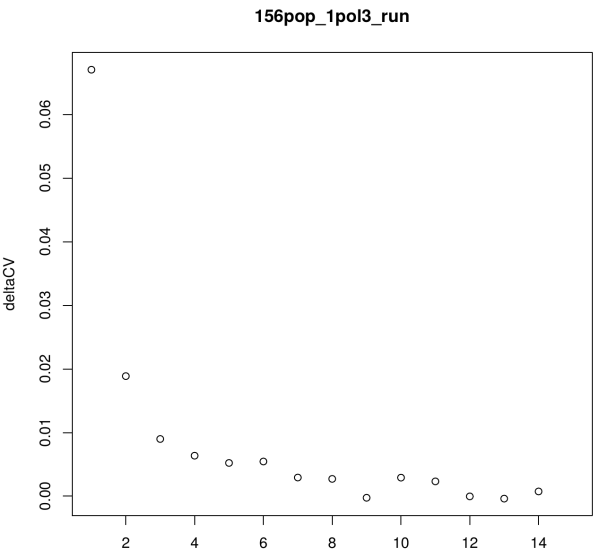

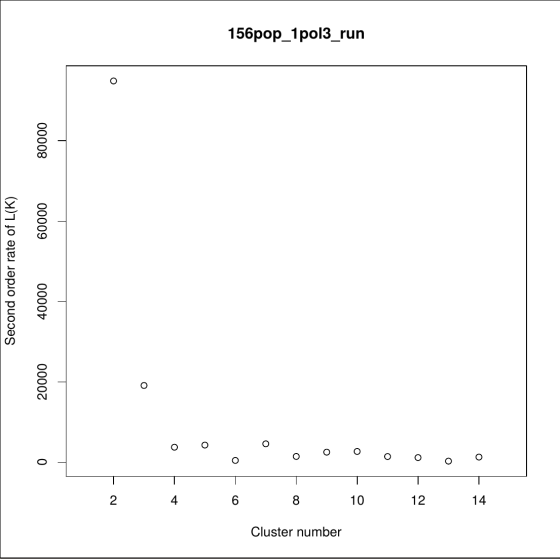

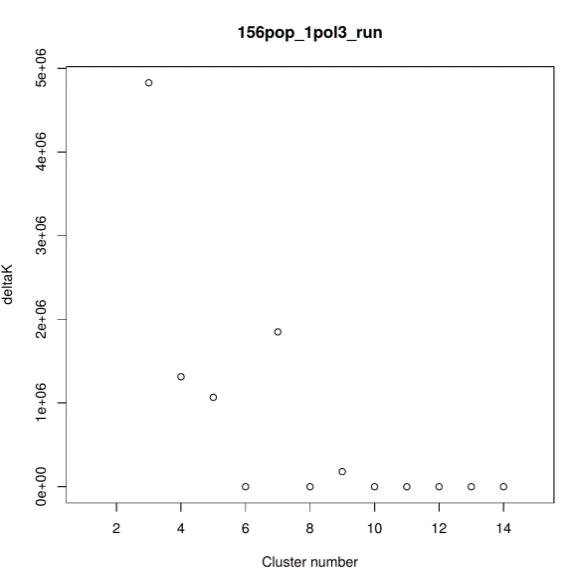

A)

B)

C)

D)
