## Supplementary material for "Genome-wide SNP genotyping of DNA pools identifies untapped landraces and genomic regions that could enrich the maize breeding pool": Fig. S8

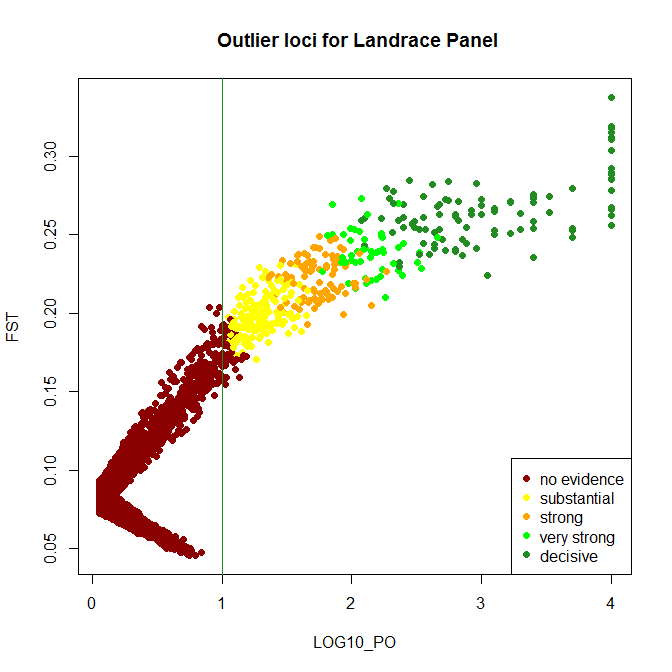

#### Fig. S8: Genomic scan performed by BAYESCAN to identify outlier loci within the landrace panel. Each point corresponds to an SNP locus. The estimated FST is plotted against the log_10_ of the posterior odds (PO), which provides evidence as to whether the locus is under selection or not. The vertical green line shows the decisive threshold value (log10_PO =1) used for identifying outlier loci.
