## Supplementary material for "Genome-wide SNP genotyping of DNA pools identifies untapped landraces and genomic regions that could enrich the maize breeding pool": Fig. S9

#### Fig. S9: Variation in the level of genetic differentiation between pairs geographic groups of landraces along the maize genome. Vertical gray bars correspond to centromere locations. Chromosome boundaries are shown as vertical dashed lines. The horizontal dashed lines correspond to the mean, 5% and the 95% percentile. Outlier regions are identified by red asterisks. The genome location of ID1, tb1, pbf1, su1, tga1, bt2, o2, pebp8, vgt1, nac1 and ZmCCT genes is shown at the top and by vertical blue lines.
