## Supplementary material for "Genome-wide SNP genotyping of DNA pools identifies untapped landraces and genomic regions that could enrich the maize breeding pool": Fig. S10

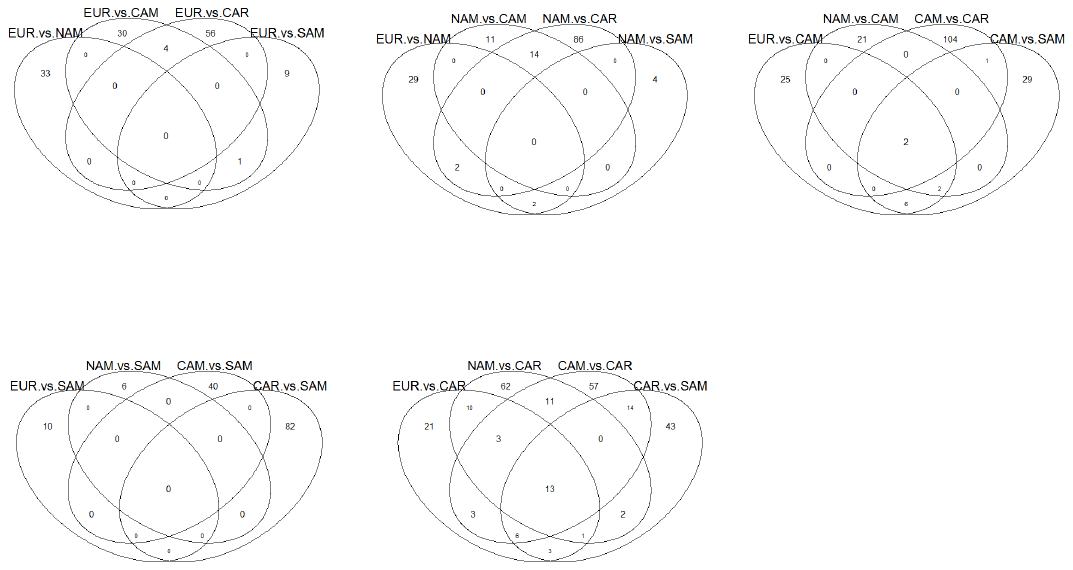

#### Fig. S10: Distribution of outlier loci in the five geographic groups of maize landraces. The Venn diagrams illustrate the number of outlier loci that are exclusive to all possible pairwise combinations among the five geographic groups of maize landraces.
