## Supplementary material for "Genome-wide SNP genotyping of DNA pools identifies untapped landraces and genomic regions that could enrich the maize breeding pool": Fig. S11

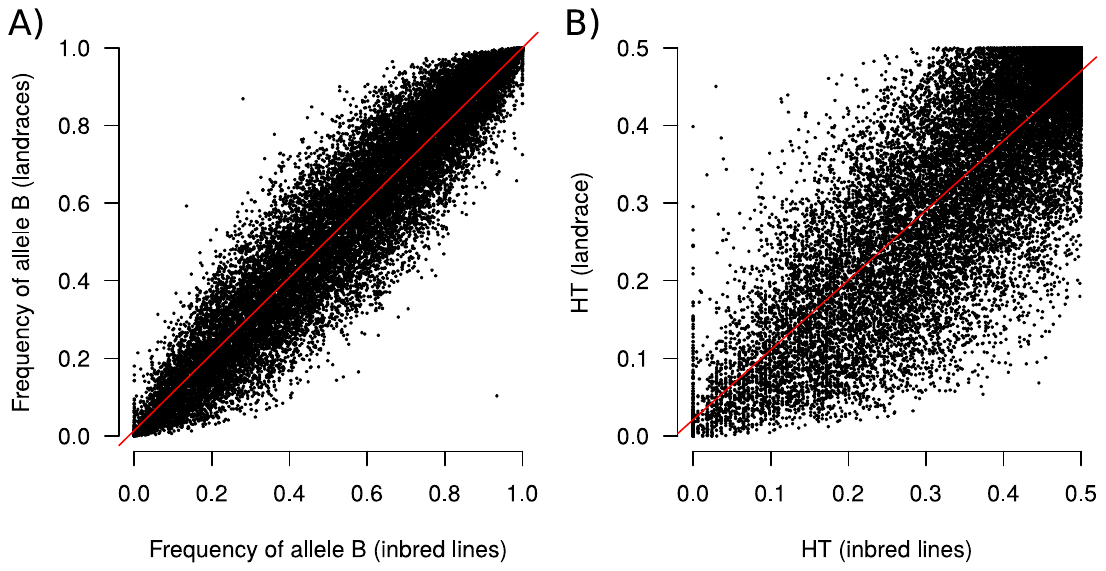

#### Fig. S11: Relationship between the mean frequency of allele B (A) and expected heterozygosity (HT, B) in panels of 166 landraces and 327 inbred lines for 23,412 SNPs. Red lines represent the linear regression. The coefficients of determination (R ^2^ ) are 0.89 and 0.71 for the frequency of allele B and HT, respectively.
