## Supplementary material for "Genome-wide SNP genotyping of DNA pools identifies untapped landraces and genomic regions that could enrich the maize breeding pool": Fig. S12

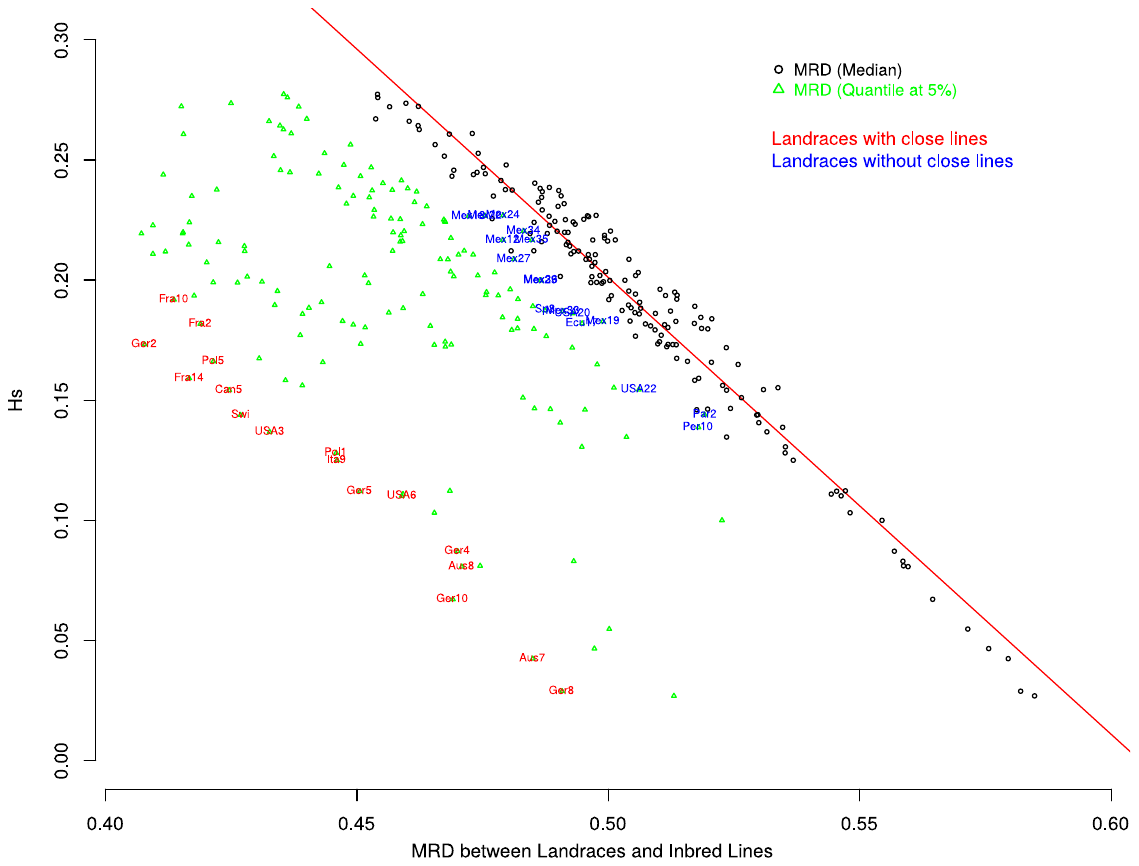

#### Fig. S12: Relationship between landrace genetic diversity (Hs) and their modified Roger’s distance (MRD), and the inbred lines from the panel of CK lines. Black circles and green triangles represent median (MRD_med_) and lower percentile (5%, MRD_05_) of MRD. The dotted red line represents the linear regression between MRD_med_ and Hs. Red and blue labels represent the 17 closest and most distant landraces to inbred lines according to the MRD normalized by Hs (MRD_norm_).
