## Supplementary material for "Genome-wide SNP genotyping of DNA pools identifies untapped landraces and genomic regions that could enrich the maize breeding pool": Fig. S13

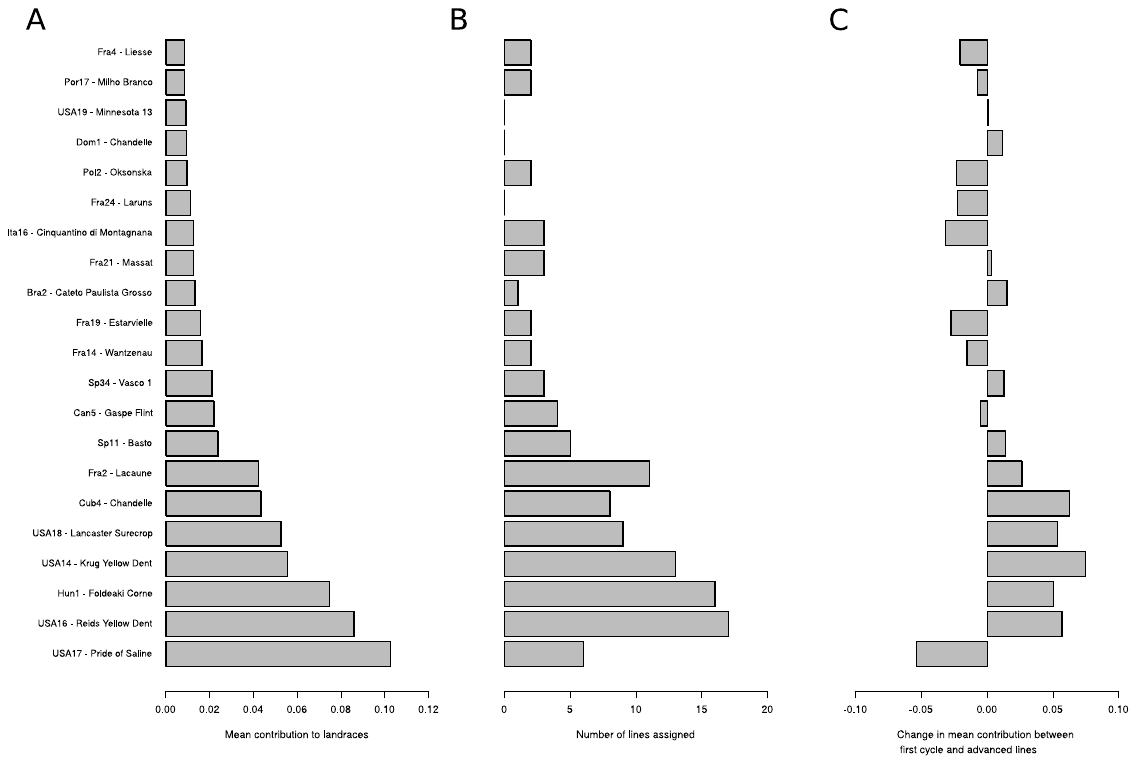

#### Fig. S13: Landraces with the highest average contribution to the 327 inbred lines from the diversity panel. (A) Average contribution of 21 landraces; (B) Number of lines assigned to these 21 landraces with an assignment probability > 60 %; (C) Change in average contribution of landraces between first cycle lines and more advanced lines from the diversity panel.
