## Supplementary Information for "Genome-wide SNP genotyping of DNA pools identifies untapped landraces and genomic regions that could enrich the maize breeding pool"

### Supplementary text 1: Selection and sampling of landraces

156 landraces were selected from 473 landraces under the following criteria: (i) maximize the genetic distance between landraces, (ii) preferentially retain landraces for which we had inbred lines derived directly by selfing (first cycle inbred lines in our “CK line panel”), and (iii) represent in equal numbers native American and European landraces. 145 populations were selected from the 275 landraces described in Camus-Kulandaivelu et *al.* (1) by applying the maximum length subtree (MLS) procedure implemented in the DARwin software (2) on the Neighbor-Joining tree of the 275 landraces built from modified Roger’s distance of 17 SSR markers. The thirty-six European and 35 American populations used to produce inbred lines were defined as forced units. We analyzed European and American landraces separately to avoid an over-representation of American landraces due to their higher genetic diversity. The MLS procedure was repeated until it reached a sample size of 72 and 73 European and American landraces, respectively. In order to enlarge the geographic coverage, we added the 11 landraces from Mir et al., (3), which were not included in Camus-Kulandaivelu et al. (1). These comprised two European landraces from Portugal, and nine American landraces originating from Argentina (2), Brazil (2), Paraguay (2), Peru (1), Uruguay (1) and Chile (1).

Each population was represented by either one (146) or two (10) DNA pools from 15 individual plants, mixed in equal amounts as described in Reif *et al.* (4) and Dubreuil *et al.* (5). In order to assess the effect of sampling on allelic frequency estimation (6), ten populations were represented by two DNA pools of 15 independently sampled plants. These populations have the same accession code and name but have different DNA sample names (Table S1).

### Supplementary text 2: Filtering SNPs according to their suitability for diversity analysis and their quality for predicting allelic frequency

50K genotyping was performed according to the manufacturer's instructions using the MaizeSNP50 array (IlluminaInc, San Diego, CA). Genotype results were produced with GenomeStudio’s Genotyping Module software (v2010.2, Illumina Inc) using the cluster file MaizeSNP50_B.egt available from Illumina. 49,585 SNPs passed the quality criteria as described in Ganal et al. (2011). We selected a subset of 30,068 SNP markers from the Panzea project (7), called prefixed PZE SNPs, which are suitable for genetic diversity analysis (8). We then used a weighted deviation criterion (wd) < 50 to identify poorly predictive SNPs for allelic frequency in DNA pools (see equation 1 in (6) for wd calculation) and discarded 6,412 SNPs. In addition, we removed 244 SNPs that were heterozygous in one of the parental lines of the controlled pools because they displayed a high rate of error in allelic frequency prediction. We obtained a final set of 23,412 SNPs for the diversity analysis.

### Supplementary text 3: Estimation of “within group” and “across group” diversity parameters and FST.

Let assume that the 166 landraces can be clustered into g groups according to their geographical origins. We can define for each SNP:

- *f_glk_* as the allelic frequency at locus k in landraces l in group g
- $\overline{f_{gk}}$ = $\frac{\sum_{l=1}^{Ng} f_{glk}}{Ng}$ as the average allelic frequency at locus k in group with Ng number of landraces in group G
- $\overline{f_{k}}$ = $\frac{\sum_{l=1}^{N} f_{glk}}{N}$ as the average allelic frequency at locus k in the whole landrace panel with N number of landraces

Considering *f_glk_* the allelic frequency of landrace l at one locus k, we can estimate the expected heterozygosity *Hs_glk_* for each landrace l at each biallelic SNP as follows:

*Hs_glk_ = 2 f_glk_ (1-f_glk_)*

A similar expected heterozygosity can be computed at the level of the group g using two different formula depending whether random mating (RM) is assumed at the level of landraces or at the level of group:

$\overline{{Hs}_{wl}}=\frac{1}{Ng}\sum_{l=1}^{Ng} 2 f_{glk} (1-f_{glk})=\frac{1}{Ng}\sum_{l=1}^{Ng} {Hs}_{glk}$ with *Ng* the number of landraces in group “g” (RM within landraces only)

$\overline{{Hs}_{wg}} =2 \overline{f_{gk}} (1-\overline{f_{gk}} )$ (RM within group)

Lastly, depending on whether random mating is assumed to occur at the landrace, group or population level, the expected heterozygocity could be defined as follows:

$\overline{{HT}_{l}}= \frac{1}{N} \sum_{l=1}^{N} 2 f_{glk} \left( 1-f_{glk} \right)$(RM within landraces only), with N number of landraces

$\overline{{HT}_{g}}= \sum_{1}^{g} \sum_{g=1}^{Ng} \frac{Ng}{N}2 \overline{f_{gk}} (1-\overline{f_{gk}} )$(RM within groups only), with N and Ng number of landraces in the whole landraces and within each group, respectively, and g number of groups

$\overline{{HT}_{p}}= 2 \overline{f_{k}} (1-\overline{f_{k}} )$ (RM in the whole population)

In Table 1 and Fig. 3, $\overline{{Hs}_{wl}}$ and $\overline{{HT}_{l}}$ corresponds to the “average of within pop expected heterozygosity Hs” whereas $\overline{{Hs}_{wg}}$ and $\overline{{HT}_{p}}$ corresponds to the “total expected heterozygosity HT” in each geographical groups and in landraces panel, respectively.

Genetic differentiation (FST) was estimated at each locus and across all loci as *per* (76, 77). The genetic differentiation between landraces among whole landrace panel (FST_l_), between 5 geographical groups among whole landraces panel (FST_g_), between landraces among each geographical groups (FST_lg_), between pairs of geographical groups (FST_pg_), between inbred lines and landraces (FST_li_) was estimated as follows (Table 1):

${FST}_{l}=1- \frac{\overline{{Hs}_{wl}}}{\overline{{HT}_{p}}}=1- \frac{\frac{1}{Ng}\sum_{l=1}^{Ng} {Hs}_{glk}}{2 \overline{f_{k}} (1-\overline{f_{k}} )}$ (1) with Ng = number of landraces

${FST}_{g}=1- \frac{\overline{{HT}_{g}}}{\overline{{HT}_{p}}}=1- \frac{\overline{\sum_{1}^{g} \sum_{g=1}^{Ng} \frac{Ng}{N}2 \overline{f_{gk}} (1-\overline{f_{gk}} )}}{2 \overline{f_{k}} (1-\overline{f_{k}} )}$ (2) with “g” representing each geographical groups, and Ng number of landraces within each geographical groups “g”.

${FST}_{lg}=1- \frac{\overline{{Hs}_{wl}}}{\overline{{Hs}_{wg}}}= 1- \frac{\frac{1}{Ng}\sum_{l=1}^{Ng} {Hs}_{glk}}{2 \overline{f_{gk}} (1-\overline{f_{gk}} )}$ (3) with Ng number of landraces within each geographical group “g”.

${FST}_{pg}=1- \frac{\overline{{HT}_{g}}}{\overline{{HT}_{p}}}=1- \frac{\overline{\sum_{1}^{g} \sum_{g=1}^{Ng} \frac{Ng}{N}2 \overline{f_{gk}} (1-\overline{f_{gk}} )}}{2 \overline{f_{k}} (1-\overline{f_{k}} )}$ (4) with “g” representing two geographical groups and assuming that whole population are the landraces originated from these two geographical groups, only.

Considering landrace and CK lines as a unique population, we can estimated FST between inbred lines and landraces (FST_li_) in the same way:

${FST}_{li}=1- \frac{\overline{{HT}_{g}}}{\overline{{HT}_{p}}}=1- \frac{\overline{\sum_{1}^{g} \sum_{g=1}^{Ng} \frac{Ng}{N}2 \overline{f_{gk}} (1-\overline{f_{gk}} )}}{2 \overline{f_{k}} (1-\overline{f_{k}} )}$ with g representing two groups: landrace panel and CK line panel and $\overline{f_{k}}$are the average frequency of locus k in whole population obtained by merging landrace and CK line panel.

The same method was applied to estimate the diversity parameters Minor Allelic Frequency (MAF) and Number of Alleles “A” across and whithin groups for geographical groups, whole landrace panels and inbred lines panel.

### Supplementary text 4: Classification of SNP by bayescan

For the BAYESCAN analysis, a Bayes factor (BF), which is the ratio of the posterior probabilities of two models (selection *vs* neutral), is estimated at each locus. According to this Bayes factor and Jeffrey’s scale of interpretation, each locus can be classified as showing substantial (log_10_(BF)>0.5), strong (log_10_(BF)>1), very strong (log_10_(BF)>1.5) or decisive (log_10_(BF)>2) evidence of selection. A log_10_(BF) of 0.5, 1, 1.5, 2 corresponds roughly to a posterior probability of 0.76, 0.91, 0.97, 0.99, respectively. Note that comparison of landraces and inbred lines at individual loci/genomic regions allowed us to identify candidate loci for genes that experienced selection during the transition to hybrid breeding.

### Supplementary text 5: Simulation of individuals for structure analysis of landraces

To simulate the genotypes of five individuals *per* population at each SNP, we sampled A and B alleles from the binomial distribution with sampling probabilities proportional to their respective frequency in the population. To avoid linkage disequilibrium-related artifacts, we analyzed a subset of 2,500 SNPs, obtained by dividing the maize genetic map into 2,500 non-overlapping windows and randomly selecting a single SNP in each window. For landraces with duplicated DNA samples, we averaged the allelic frequencies of duplicated DNA samples, with the exception of Pol3 (a landrace from Poland) for which duplicated samples were highly divergent (6). Based on hierarchical clustering analysis (Figure 1B) we included the Pol3 DNA sample DivCor_137b, which was close to Pol2 (another landrace from Poland) and excluded the Pol3 DNA sample DivCor_137a, which was close to Chi13 (a landrace from Chile).

The robustness of ADMIXTURE assignments using these 2,500 SNPs was tested by comparing these with the cluster assignments obtained using STRUCTURE (9) on 17 SSR markers previously analyzed by Mir et al., (3). To check for agreement between the results of the two assignment methods, we plotted the relationship between Q group membership coefficients and estimated Pearson's correlation coefficient, as in Hamblin et al. 2007 (Table S2).

1.	L. Camus-Kulandaivelu, Maize Adaptation to Temperate Climate: Relationship Between Population Structure and Polymorphism in the Dwarf8 Gene. *Genetics* **172**, 2449–2463 (2006).

2.	X. Perrier, J. P. Jacquemoud-Collet, *DARwin software* (2006).
